# Benchmarking the quantitative performance of metabarcoding and shotgun sequencing using mock communities of marine nematodes

**DOI:** 10.64898/2026.02.09.704827

**Authors:** Dandan Izabel-Shen, Henrik Sandberg, Mohammed Ahmed, Elias Broman, Oleksandr Holovachov, Francisco J. A. Nascimento

## Abstract

High-throughput sequencing has transformed biodiversity assessment and ecological monitoring, yet its quantitative reliability remains unclear. Here, we assembled two experiments of nematode mock communities: one based on extracted DNA and one on individual specimens. Although DNA extraction was required in both experiments to assess the quantitative performance of the sequencing approaches, we essentially evaluated whether this performance was influenced by differences in the start material used to constructed the mock communities. Each community was analyzed using 18S and 28S metabarcoding and shotgun sequencing to evaluate their ability to resolve quantitative information. Across datasets, the number of observed taxa increased with sequencing depth despite controlled input, indicating that higher read numbers primarily revealed intragenomic variation in nematodes than true diversity. Community composition was more accurately recovered by 18S metabarcoding and shotgun sequencing than by 28S. Both sequencing approaches reflected DNA input reasonably well; however, shotgun sequencing provided more consistent abundance estimates relative to individual counts, particularly for nematodes with relatively large-bodied size. In contrast, all methods showed limited ability to accurately quantify taxa with low DNA input or small body size. Comparisons between mock community types showed strong correspondence between read abundance and DNA input, but weaker relationships with individual counts. Overall, both metabarcoding and shotgun sequencing effectively detected community-level patterns and within-taxon abundance, but shotgun sequencing was more reliable for cross-taxon quantitative comparisons. Our findings demonstrate how input material, primer choice, and sequencing approach influence the accuracy of nematode abundance estimates, and provide guidance for improving quantitative applications in nematode-based bioindication and, more broadly environmental DNA biomonitoring.

## Introduction

High-throughput sequencing has advanced our awareness of both the visible and the previously invisible biodiversity (Bik et al., 2012; Cook et al., 2025). Among the novel applications of DNA metabarcoding are the creation of reference databases across the Tree of Life (Cook et al., 2025; Holovachov, 2016) and as a cost-efficient tool for the biomonitoring of aquatic organisms (Fonseca et al., 2010). In benthic environments the meiofaunal phylum Nematoda is highly abundant, diverse, and plays an important role in food web dynamics and biogeochemical processes in the sediment (Bonaglia et al., 2014; Nascimento et al., 2012; Schratzberger and Ingels, 2018; van Der Heijden et al., 2020). With their rapid growth and short-life cycle, nematodes can rapidly respond to environmental disturbances, making them valuable bioindicators for detecting early signs of ecosystem changes (di Montanara et al., 2022; Ridall and Ingels, 2021). However, traditional morphology-based methods of nematode identification are laborious and require well-trained taxonomists, thus the utility of nematodes in biomonitoring has been limited (Kennedy and Jacoby, 1999). With the advent of DNA metabarcoding analysis, this and other limitations of meiofauna-based biomonitoring have been circumvented (Creer et al., 2010; Fonseca et al., 2010). Consequently, research on nematode-based indicators of environmental impact has increased substantially in recent years (di Montanara et al., 2022). There is also a growing number of studies using DNA metabarcoding to examine relationships between meiofaunal community dynamics and environment factors across large spatial scales (e.g., Broman et al., 2019; Frontalini et al., 2025; Iburg et al., 2021).

The accurate quantification of biological communities relies on detection, abundance estimates, and taxonomic representation of individual organisms in that communities. This knowledge also underpins most benthic monitoring indices, which are based not only on taxon presence or absence but also on their relative abundances (Pinto et al., 2009). As such, metabarcoding approaches for detecting ecological patterns and quantifying taxon abundance face two major challenges (Lamb et al., 2019; van der Loos and Nijland, 2021). First, there is substantial variation both within and between taxa in key genome attributes, such as nucleotide sequence, cell number, and target gene copy number; such variation is especially pronounced among eukaryotes (Bik et al., 2012). Second, the inherent biases introduced during PCR amplification can lead to over- or under-representation of certain taxa in the sequence dataset relative to their true abundances in natural communities (Piñol et al., 2015). These issues can distort ecological patterns in biological communities when sequences and read counts are used to assess taxonomy and abundance, hindering reliable assessments of their ecological status. Mock community experiments, like the present study, where known materials from various taxa are combined, have been designed to benchmark sequencing biases and evaluate the effectiveness of different methods in quantifying the relative abundance or biomass of included organisms.

Since larger individuals typically contain more DNA, sequence read counts tend to correlate more closely with relative biomass than with relative abundance based on individual counts. However, the degree of the association between the proportion of reads and either abundance or biomass varies considerably across different studies. A meta-analysis of metabarcoding mock community experiments by Lamb et al., (2019) found a significant but weak positive correlation between the proportion of reads and the amount of input material. They found no consistent difference whether input was measured as number of individuals, total biomass, or DNA/RNA quantity. Additionally, they observed that in studies using individual counts as the input measure, the individuals were likely of similar size. Other reviews have corroborated these findings, showing that correlations between sequencing reads and input material vary widely across studies, ranging from strong to non-existent (Deagle et al., 2019; van der Loos and Nijland, 2021). Consequently, the quantitative reliability of metabarcoding must be evaluated on a case-by-case basis, as it depends on the target community and the sequencing approach used. For meiofauna, Ershova et al., (2021) reported a significant correlation between read counts and biomass proportions in a study of marine zooplankton using the COI marker. In nematodes, Schenk et al., (2019) observed significant correlations between read proportions and both abundance and biomass, with a stronger relationship observed for biomass. These findings were consistent when using primers targeting either the D3 region of 28S or the V4 region of 18S. However, when primers targeting the V1 region of 18S were employed, the results were highly skewed, showing a pronounced overrepresentation of a single organism. These findings highlight that metabarcoding performance is likely primer-specific, and results often not generalizable across different target genes or even regions within the same gene. Therefore, additional research is necessary before definitive conclusions can be made regarding the most effective metabarcoding approaches for meiofaunal communities.

Shotgun sequencing offers an alternative method to investigate the structure and composition of biological assemblages, as its sequences all random DNA within a community sample, eliminating the need for PCR amplification of a marker gene. However, this method has drawbacks, including high costs and reliance on the availability and quality of reference genomes for genome assembly and taxonomic annotation (Elliott and Coissac, 2025; Nasko et al., 2018). Despite these challenges, shotgun sequencing has shown potential in estimating the biomass of various organism groups, such as benthic macrofauna (Bista et al., 2018; Callens et al., 2025) and soil invertebrates (Schmidt et al., 2022). Furthermore, a previous study by (Paula et al., 2022) demonstrated that shotgun sequencing achieved consistent quantitative performance. Similar to metabarcoding, there is a need for more systematic benchmarking studies to evaluate the effectiveness of shotgun sequencing for meiofauna in general and nematodes in particular. Beyond biases inherent in the sequencing process, the completeness of reference databases and the representation of targeted taxa within them are crucial factors influencing quantitative accuracy Genetic studies of free-living meiofauna commonly utilize the 18S rRNA gene, often in conjunction with the 28S rRNA gene. Both markers are highly conserved, enabling the targeting of a wide range of taxa (Gielings et al., 2021; Schenk and Fontaneto, 2020; van der Loos and Nijland, 2021). Historically, well-used marker genes and regions such as the 18S and 28S rRNA genes are better represented in references databases, whereas the completeness of those databases for other markers may be inadequate (Holovachov et al., 2017). Although shotgun sequencing is a PCR-free approach, it still relies on reference databases for taxonomic classification, making its accuracy contingent on database quality. Additionally, shotgun sequencing is currently less cost-effective than metabarcoding, which limits its use in large-scale meiofaunal community studies.

As noted earlier, best practices for sequencing studies of meiofauna, especially nematodes, remain to be established, as they depend on understanding of the factors that influence the relationship between read abundance and biomass or individual counts (Lamb et al., 2019). Both PCR-based and PCR-free methods may contain systematic biases, which should be thoroughly examined through direct comparisons. Identified biases can potentially be corrected during data analysis, for example, by using correction factors from benchmarking studies (Thomas et al., 2016). Therefore, experiments to evaluate the quantitative accuracy of various sequencing methods are essential for developing reliable protocols.

In this study, mock communities consisting of eight Baltic Sea nematode genera were assembled to evaluate the quantitative reliability of DNA metabarcoding and shotgun sequencing. This design enabled the comparison of organism detection, taxonomic assignment, occurrence estimates, and abundance quantification across sequencing approaches. These genera are abundant in the Baltic Sea (Broman et al., 2019) and differ in their sensitivity to hypoxia (Modig and Ólafsson, 1998), making them suitable indicators of both persistent hypoxia in deeper basins (Kuliński et al., 2022) and seasonal hypoxia in coastal areas (Conley et al., 2011). Mock communities comprising either DNA extracted from nematodes or whole nematode specimens were established to assess how sequencing read abundances reflect expected proportions based on DNA quantity or individual counts, respectively. Using DNA as input in the mock assembly allowed us to mitigate DNA extraction biases associated with organismal traits, such as body size and cuticle thickness in nematodes, which are difficult to control when extracting DNA from multiple genera simultaneously. The study aimed to test three hypotheses: (1) that metabarcoding and shotgun sequencing produce similar community diversity patterns reflecting the mock community composition; (2) that relative abundances from sequencing data correlate more strongly with the input DNA quantity for each genus, with shotgun sequencing expected to show a higher correlation than metabarcoding; (3) that shotgun data more accurately reflect the number of individuals in the mock communities than metabarcoding. By identifying potential methodological biases and the need for correction factors, our findings contribute to the development of optimized protocols for accurate, sequencing-based quantification of nematode communities, supporting the use of these organisms in environmental monitoring programs.

## Materials and Methods

### Sediment collection

The top 2 cm of sediment was collected from multiple stations of the Baltic Proper during four consecutive years (2016–2019) and from the Bothnian Sea and the Bothnian Bay as previously reported (Broman et al., 2022, 2019). The sieving methods for the sediments and the isolation of meiofauna from sites where nematodes were the most abundant component were previously described (Broman et al., 2019). Briefly, sediment samples were sieved through a sterilized 40-µm sieve (autoclaved, rinsed with MilliQ water between samples). Organisms in the sieved sediment retained on the 40-µm sieve were isolated by density extraction using a Levasil silica gel colloidal dispersion solution. The top part of the solution, containing the meiofauna, was decanted and washed with distilled water. This isolation procedure was repeated twice and the pooled content of the three isolations was placed in a 40-µm sieve and washed thoroughly with distilled water to remove residual Levasil. The 40-µm sieve content, containing the meiofauna isolate, was transferred into a 15-mL sterile Falcon tube with a maximum final volume of 10 mL. All meiofauna suspensions were frozen at –20°C until microscopic identification of nematodes for the experiments.

### Nematode sorting and identification

Before microscopic identification and nematode sorting, these meiofauna suspensions were thawed, and the nematode suspension was concentrated using a 46-µm sieve. Aliquots were examined in counting dishes, and nematodes were sorted under an Olympus SZX12 stereo microscope equipped with Olympus DF PLFL 1.6 X PF objective, and WHS10X/22 eyepieces at the highest magnification. Subsequently, individual nematodes were morphologically identified using a Nikon Eclipse 80i DIC microscope with cavity glass slides. Specimens were identified at the genus level and assigned to the eight target nematode genera used for the mock community assembly: *Sabatieria*, *Halomonhystera, Leptolaimus*, *Paracanthonchus*, *Microlaimus*, *Daptonema*, *Axonolaimus*, and *Desmolaimus*. Ten individuals of the same genus were transferred to a sterile Epi-tube filled ultra-clean nucleic acid-free water (Thermo Fisher Scientific), and stored at –20°C again (**Figure 2**) until use in the mock community experiments.

Given the taxonomic and morphological diversity within Nematoda, we consider genus-level identification of target nematodes in our mock community design to be a practical and robust compromise. This approach minimizes ambiguity in taxonomic assignment and reflects realistic research scenarios.

### Mock community design

Three ecological groups of nematodes were classified according to their sensitivity to hypoxia (Modig and Ólafsson, 1998) (**Table 1 and Figure 1**): i) *Sabatieria*, *Leptolaimus*, *Halomonhystera* (hypoxia-tolerant), ii) *Paracanthonchus*, *Daptonema*, *Microlaimus* (less hypoxia-sensitive), iii) *Axonolaimus*, *Desmolaimus* (hypoxia-sensitive). Prior to assembling the mock communities composed of DNA input, genomic DNA was extracted from morphologically identified nematodes stored in Epi-tubes (**Figure 2**). Multiple tubes representing the same genus were processed to obtain sufficient DNA from each genus for the construction of the mock communities. Three types of staggered mock communities were assembled in Experiment 1, each differing in the relative contribution of ecological groups and replicated three times (n=9; **Figure 3A**): (i) communities dominated by DNA from tolerant taxa; (ii) communities dominated by DNA from less-sensitive taxa and (iii) communities dominated by DNA from sensitive taxa. Although the DNA quantity of the different ecological groups differed within each mock community (**Supplementary Table S1**), the total DNA yields were kept the same, with 22 ng DNA for each community. Experiment 2 included four types of mock communities (three replicates each; n=12): one with equal numbers of individuals per target genus, and three with unequal numbers representing dominance of tolerant, less-sensitive, or sensitive taxa, correspondingly (**Figure 3B**). Each assembly consisted of a total of 80 individual nematodes but the number of individuals per genus varied, except in the case of the even community, which had a total of 70 individuals because *Daptonema* had to be excluded due to a lack of specimens.

**Table 1.** Selection of Nematoda genera differing in their sensitivity to hypoxia in the Baltic Sea.

| Nematoda | Body size | Hypoxia sensitivity |
| --- | --- | --- |
| *Sabatieria* | intermediate thick-bodied | Tolerant |
| *Halomonhystera* | large | Tolerant |
| *Leptolaimus* | Small | Tolerant |
| *Paracanthonchus* | intermediate slender-bodied | Less sensitive |
| *Microlaimus* | Small | Less sensitive |
| *Daptonema* | intermediate thick-bodied | Less sensitive |
| *Axonolaimus* | large | Sensitive |
| *Desmolaimus* | intermediate slender-bodied | Sensitive |

**Figure 1.**
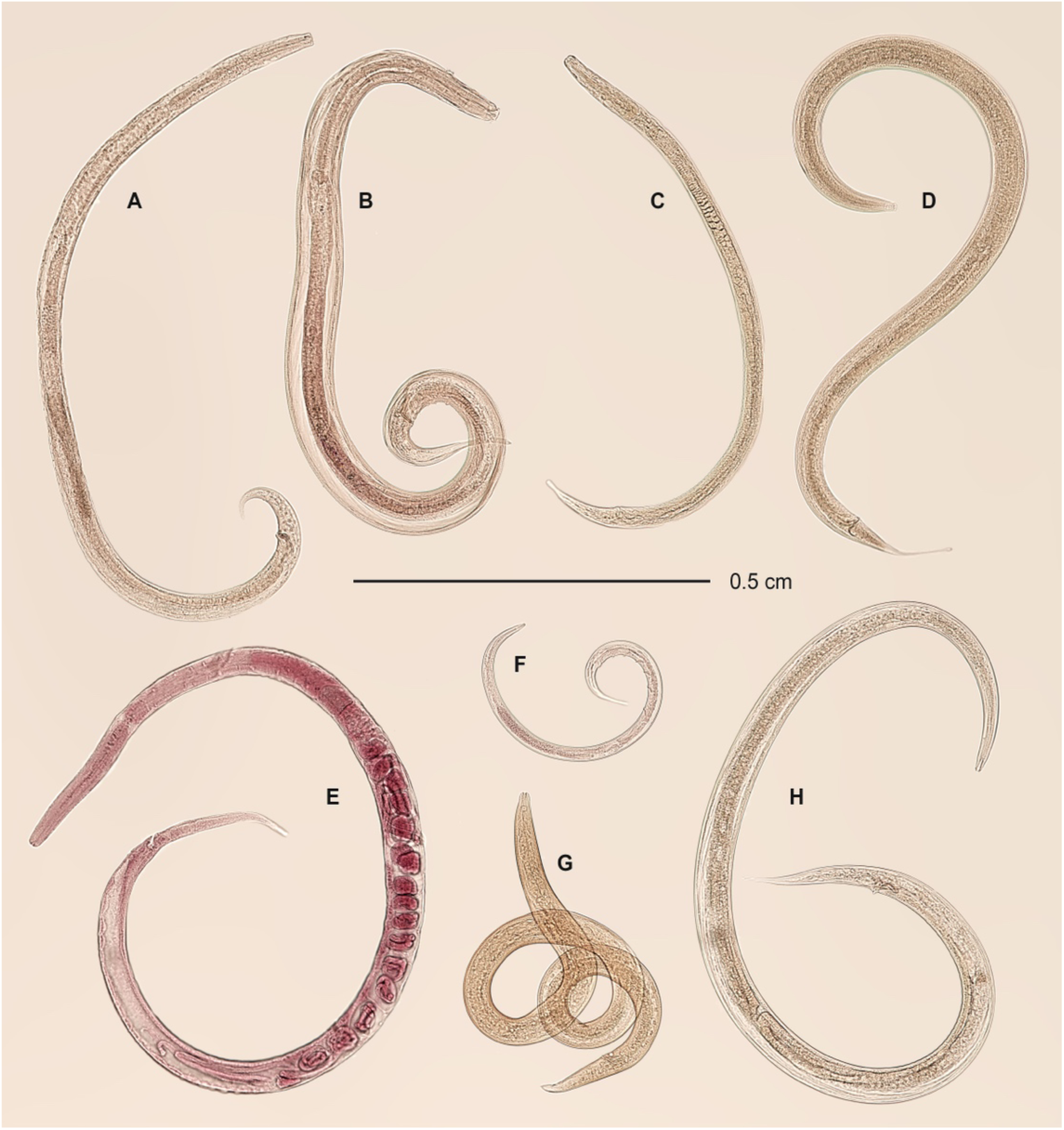
Microscopic images of the nematode genera examined in this study. A: *Paracanthonchus*; B: *Daptonema*; C: *Desmolaimus*; D: *Sabatieria*; E: *Halomonhystera*; F: *Leptolaimus*, G: *Microlaimus*; H: *Axonolaimus*.

**Figure 2.**
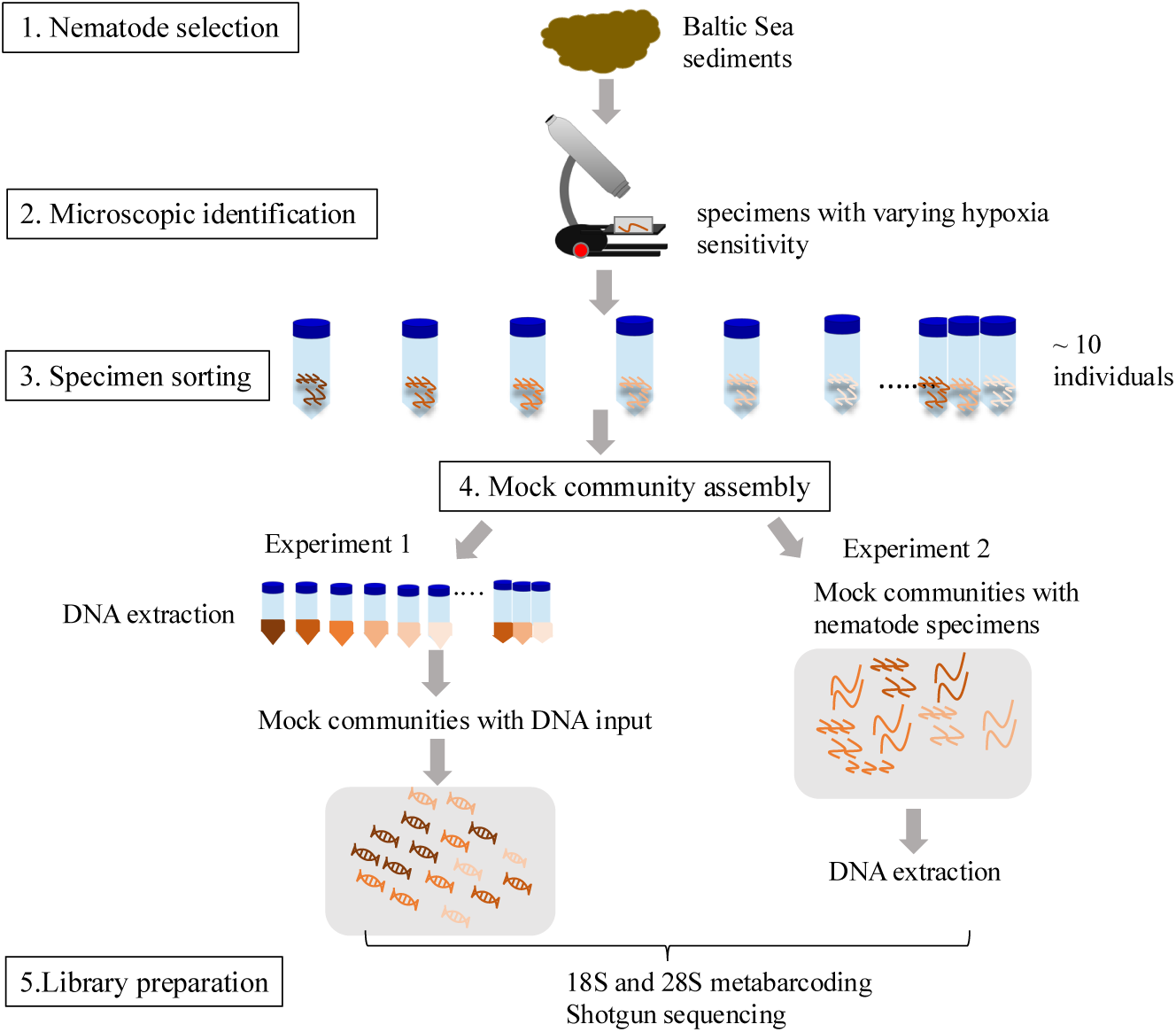
Workflow of the mock community experiment: nematode sampling from sediments of the Baltic Sea, specimen selection via microscopic identification, construction of the mock communities, and sequencing. For Experiment 1, DNA was extracted from separate tubes, each containing 10 individuals of the same morphologically identified genus to ensure efficient lysis. DNA from each extraction was used for a single replicate only and was not shared across replicates.

**Figure 3.**
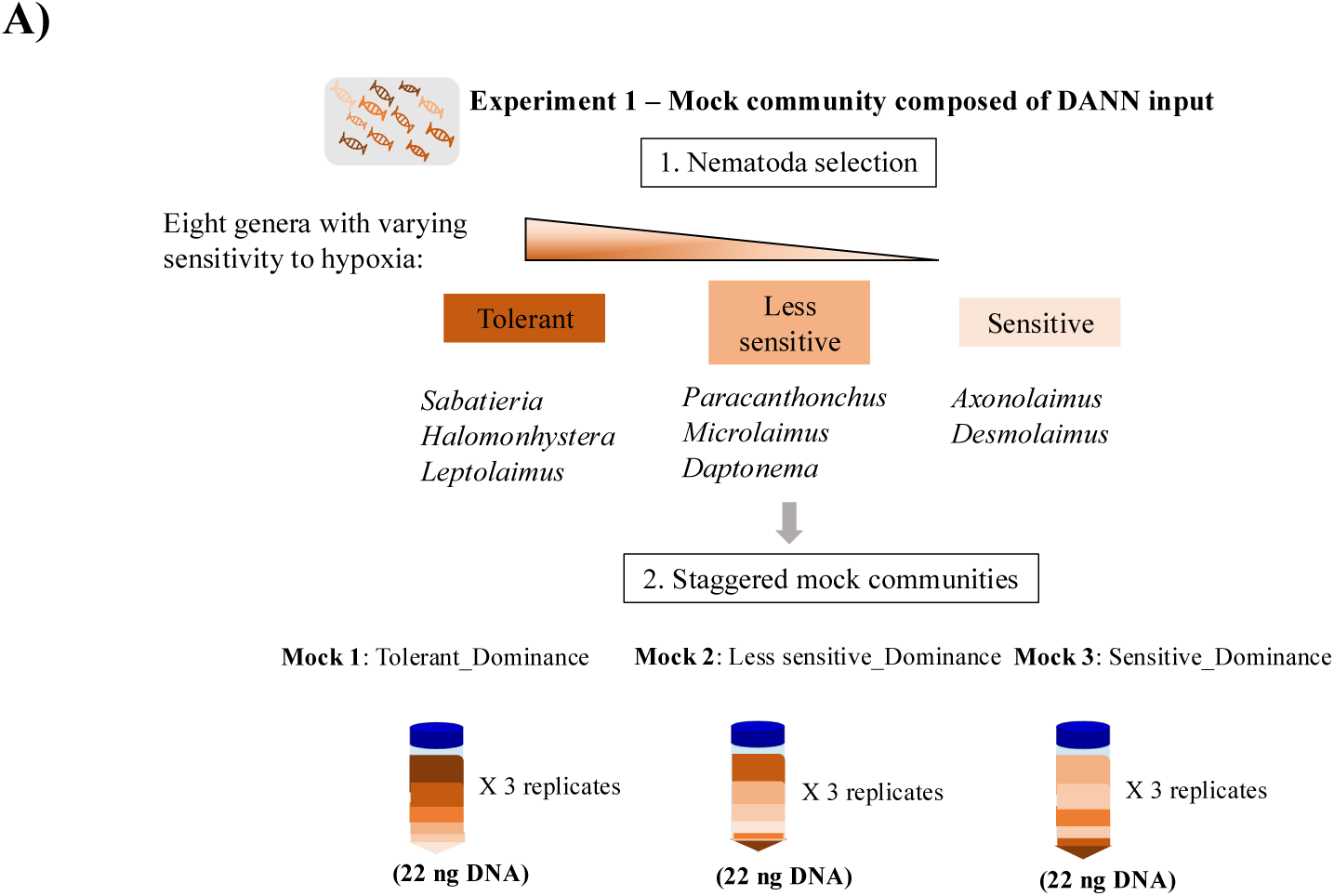
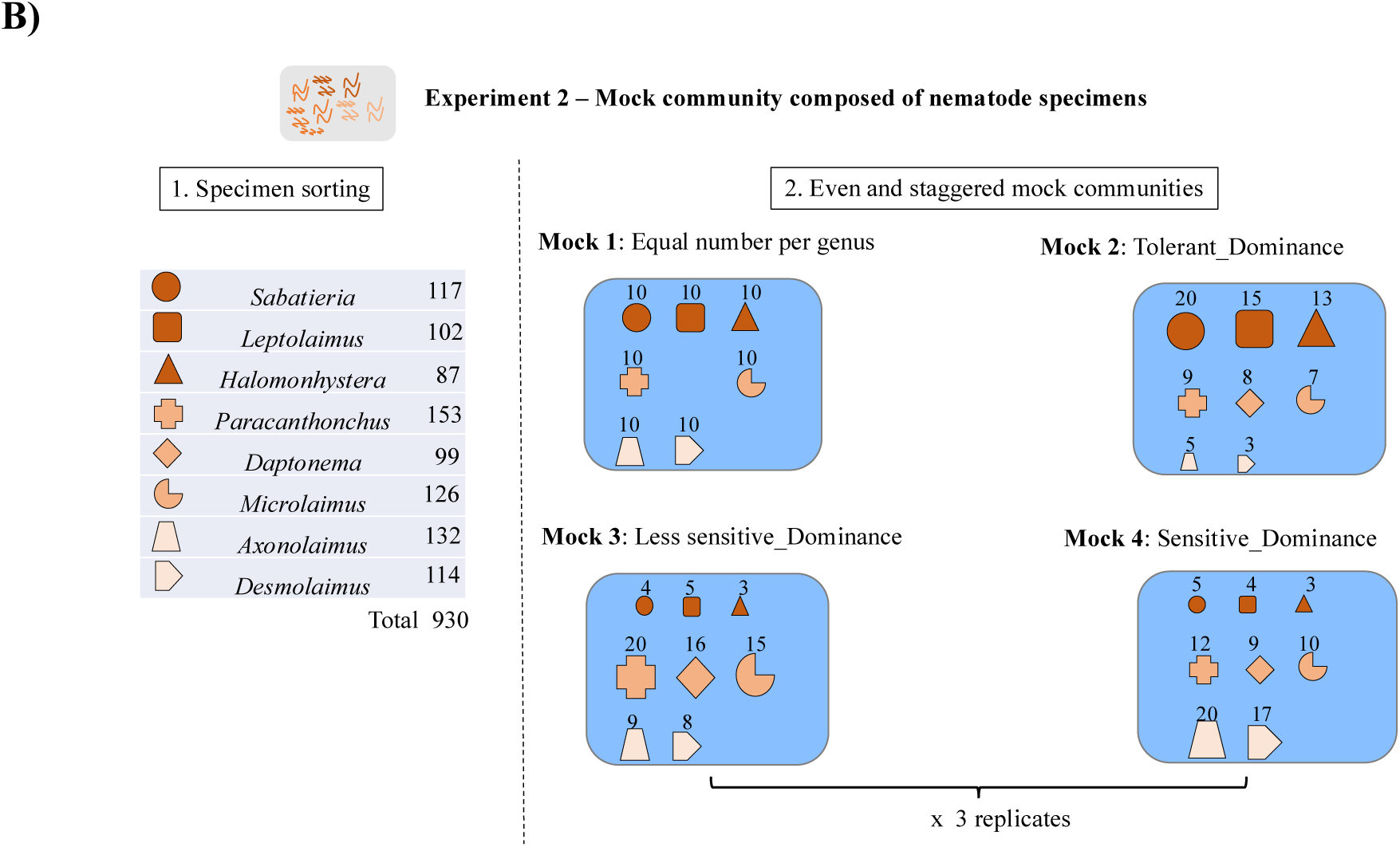
Experimental design used in mock community construction. **A**) Experiment 1: three mock community types were established by pooling different DNA quantities extracted from specific nematode taxa. For example, in “Mock 1 Tolerant_Dominated” the amount of DNA from hypoxia-tolerant genera was larger than that from genera less-sensitive and sensitive to hypoxia. Details of the amount of input DNA for each Nematoda genus are provided in **Supplementary Table S1**. **B**) Experiment 2: mock community composed by pooling individual nematode specimens prior to DNA extraction, including one even community and three staggered communities. Each symbol represents one nematode genus. The values on the left of the dashed line represent the number of individuals from a particular genus included in the construction of the mock communities. The value above each symbol indicates the number of individual specimens required for the respective mock community. Four types of mock community were assembled. For example, “Mock 1 Equal number per genus” indicates that the number of individuals was the same for each nematode genus in this community, and “Mock 2: Tolerant_Dominated” that the number of individuals in the tolerant group was greater than that in the two other ecological groups. Note that *Daptonema* was excluded in the even mock community due to insufficient specimens. For both panels, color gradients represent the sensitivity to hypoxia, with dark orange indicating hypoxia-tolerant genera, medium orange genera less sensitive to hypoxia, and light orange genera with a relatively high sensitivity to hypoxia.

### DNA extraction

For Experiment 1, one DNA extraction unit contained 50 nematodes of the same genus selected from five separate epi-tubes (10 in each tube) and pooled in one 1.5-ml epi-tube. DNA was extracted from the epi-tube sample using the QIAamp DNA Micro kit (QIAGEN) following the manufacturer’s instructions, with minor modifications. Several pre-PCR tests were run to identify the DNA extraction protocol best suited for our samples. As such, RNA carrier was not included in the extraction because our pretests showed that its addition did not enhance the DNA yields. Samples treated with proteinase K (QIAGEN) were incubated overnight at 56°C on an Eppendorf Thermomixer with 375 rpm to optimize the cell lysis. The same protocol was applied for the DNA extraction from the pooled nematode specimens for Experiment 2. As a control, one blank extraction was carried out in parallel with each set of samples and during library preparation for sequencing. In total, approximately 2,800 specimens were individually identified under the microscope and used during the experimental preparation and setup, including preliminary tests to optimize cell lysis, DNA extraction, PCR amplification, and mock community assembly

### Library preparation and sequencing

Several primer sets differing in their target coverage for taxon determinations have been used in studies of nematode metabarcoding. In this study the suitability of two primer sets for use in our mock communities was examined: (1) TAReuk454FWD1 (5′-CCAGCA(G/C)C(C/T)GCGGTAATTCC-3′) and TAReukREV3 (5′- ACTTTCGTTCTTGAT(C/T)(A/G)A-3′) (Stoeck et al., 2010), to target the V4 region of 18S rRNA gene, and (2) 1274F (5′-GACCCGTCTTGAAACACGGA-3′) and 706R (5′- GCCAGTTCTGCTTACC-3′) (Markmann and Tautz, 2005), to target the D3-D5 region of 28S rRNA gene. For each step in the library preparation (DNA extraction to amplification), a negative control was included. The negative controls were analyzed with Qubit and gel electrophoresis after full library preparation and did not produce observable bands. The details of PCR amplification conditions of 18S and 28S metabarcoding are provided in **Supplementary Text 1**. Finally, DNA from each mock community sample was subjected to three types of sequencing: 18S and 28S metabarcoding as well as shotgun sequencing. Metabarcoding was performed on an Illumina MiSeq V3 platform using a 2 × 300-bp paired-end setup at the National Genomics Infrastructure (NGI), SciLifeLab, Stockholm, Sweden, whereas shotgun sequencing was performed on a NovaSeq platform at the same center.

### Metabarcoding processing

The primers and adapters of the 18S and 28S metabarcoding sequences were trimmed using cutadapt (v1.17) (Martin, 2011). Subsequently, quality control and filtering, error rate modeling, denoising, ribosomal sequence variant inference, merging and chimera removal were carried out using the DADA2 R package (Callahan et al., 2016). The 18S sequences were quality filtered as follows: truncLen = c(275, 225), maxEE = 3, truncQ = 2, maxN = 0, rm.phix = TRUE). For the 28S sequences, the quality filtering setting included: truncLen = c(290, 260), maxEE = 4, truncQ = 2, maxN = 0, rm.phix = TRUE).

### Shotgun sequencing bioinformatics

Illumina adapter sequences were removed from the raw reads using using SeqPrep v 1.2 (St. John, 2011). PhiX sequences were removed with bowtie2 v2.3.5.1 (Langmead and Salzberg, 2012) using a PhiX genome (NCBI accession: NC_001422.1) built into a bowtie2 index. Trimmomatic v0.39 (Bolger et al., 2014) were used to trim reads (using parameters: leading=20, trailing=20, min length=75) to remove low-quality bases at the start and end of each read as well as remove very short sequences. Quality of the trimmed reads was assessed using FastQC v0.11.9 (Andrews, 2010) and MultiQC v1.11 (Ewels et al., 2016) to verify the absence of adapter contamination and to confirm per-base sequence quality. To facilitate taxonomic profiling of eukaryotic rRNA sequences, the trimmed reads were first screened with SortMeRNA v4.3.4 (Kopylova et al., 2012) to extract 18S rRNA reads in the metagenomic dataset (using the default SortMeRNA database but with non SSU sequences removed). The full read set was then co-assembled using MEGAHIT v1.2.9 (Li et al., 2015) with default parameters. Co-assembly was chosen to maximize contig continuity and detection, acknowledging that single-sample assemblies may reduce cross-sample attribution. The resulting contigs were indexed and reads were mapped back to the assembly using Bowtie2 v2.4.5 (Langmead and Salzberg, 2012) with sensitive-local alignment settings. Samtools v1.14 was used to convert the .sam output files from Bowtie2 into .bam files. Count tables of mapped reads per contig per sample were generated with FeatureCounts v2.1.1 (Liao et al., 2014), with Default values aside from -p – countReadPairs to treat the data as paired-ended, -C to exclude chimeric fragments, and -f to get read count per contig. The script blastoutput2gff.sh (Kiu, 2017) was used to convert the top hit BLAST output file from the taxonomic assignment, described below, to a .gff file used as input to FeatureCount together with the .bam.

Extraction of 18S reads from the shotgun dataset provided independent validation of the 18S metabarcoding results and enabled more reliable quantification with reduced influence of marker specific PCR bias. Although 28S reads were also extracted from the shotgun data, the resulting 28S contigs were excluded from downstream analyses because their taxonomic assignments were not sufficiently informative and reliable.

### Taxonomic assignment

Taxonomic assignment was performed for all three datasets using BLAST v2.17.0+ with default settings, except that the initial E-value threshold was set to 0.001. More stringent filtering based on e-value and other BLAST statistics was applied downstream. Amplicon sequence variants (ASVs) from the 18S and 28S datasets, as well as contigs from the shotgun dataset, were used as query sequences. The 18S ASVs and shotgun contigs were queried against the NCBI core nucleotide BLAST database (core_nt, downloaded on August 15, 2025. Because 28S metabarcoding reference data for the target nematode genera were limited in public NCBI databases, two to five individuals from each nematode genus of interest were barcoded and sequenced to generate additional 28S reference sequences. Based on the phylogenetic tree of the 28S rRNA gene reference sequences, hypoxia sensitivity appeared to be a relatively conserved trait among the eight target nematode genera (**Supplementary Figure S1**). Although some genera were represented by multiple species, these species were ecologically similar in their sensitivity to hypoxia. This pattern may be even more conserved when inferred from the 18S rRNA gene, given that 18S generally contains more conserved regions. These in house reference sequences were added to the NCBI LSU reference database (LSU_eukaryote_rRNA, downloaded on May 15, 2026), to generate a custom database for BLAST searches of the 28S ASVs. For each query sequence, the top hit, based on default blastn sorting, was retained after excluding hits and did not allow genus-level assignment. Excluded hits included, for example, entries classified only as “uncultured eukaryote” or “Nematoda sp.”, as well as hits associated with multiple taxonomic IDs that differed in genus-level classification. For the shotgun dataset, the BLAST output reduced to one hit per contig was used as input for featureCounts, as described above. For final taxonomic assignment, the top hits were subjected to additional filtering. Hits were removed if they had a bit-score below 155, percent identity below 85%, query coverage below 70%, or an e-value above 1e-50. For the 18S and shotgun datasets, hits were further filtered to retain only those with “18S” or “small subunit ribosomal” in the subject title (salltitles), ensuring that only 18S-derived matches were included. Taxonomic names were assigned by matching the subject taxonomic IDs (staxids) to their corresponding lineages in the NCBI taxonomy database, based on taxdump downloaded on May 7, 2026. When a hit was associated with multiple taxonomic IDs, the first ID was retained only if all IDs had the same genus-level classification. Taxonomic lineages were subsequently cleaned to remove non-taxonomically informative entries, and the major taxonomic ranks were extracted. The resulting lineage information and BLAST statistics were then joined with the read abundance tables for each dataset.

Before downstream analyses, sequences that did not pass the blast filtering criteria were excluded, as well as singletons defined as sequences represented by only a single read within the respective datasets. In addition, sequences not assigned to the Phylum Nematoda were excluded from most analyses. However, these non-nematode sequences were retained for the analysis comparing the proportion and composition of non-target reads and are hereafter referred as “Non nematodes”.

### Statistics

Community richness and sequencing effort for each community were assessed using linear regression. The resulting R^2^ and the *p*-value of the slope were used to evaluate the strength and significance of the relationship between the number of ASVs or contigs and the number of reads per sample. Bray-Curtis dissimilarity was used to calculate the dissimilarity between each pair of communities, with mock composition estimated from relative read abundances and compared to the expected proportions based on either DNA quantity or nematode individual counts across sequencing methods. A Wilcox test was then applied to assess whether the observed differences in community composition were statistically significant. The variability of the community composition within each type of mock assemblage (DNA input or nematode specimens) was assessed in a principal coordinates analysis (PCoA) with the Bray-Curtis dissimilarity metric provided in the vegan package. A Pearson correlation analysis was performed to test the relationship between ASV relative abundances and the expected proportion of each genus of genus in the respective mock community. Thus, ASV relative abundances were correlated against DNA quantity within the mock communities composed of DNA input, and the expected proportion of the total number of nematodes within the mock communities composed of nematode specimens. The resulting coefficients and 95% confidence intervals were used to visualize the relationships between the observed relative abundances and the expected values for the 18S, 28S, and shotgun datasets.

In this study, false positives were defined as 18S ASVs and contigs that passed the BLAST filtering criteria but were assigned to nematodes outside of the target genera in the mock communities (hereafter referred to as “Other nematodes”). Wilcoxon test was used separately to assess differences in the relative read abundances of “Other nematodes” and “Non-nematodes” among the three sequencing methods. All ecological statistics and graphs were performed in R (R Core Team, 2021). False negatives were defined as the target genera in the mock samples but was not detected in the sequencing output.

## Results

### Sequencing summary and richness recovery

The number of reads assigned to nematodes retained per sample after DADA2 quality filtering, chimeric removal, and singleton exclusion ranged from 173,409 to 559, 286 in the 18S dataset, and from 18, 645 to 204, 066 in the 28S (**Figure 4**). Following bioinformatic processing of the shotgun data, a total of 262 non-singleton nematode contigs were recovered, with the number of reads per sample ranging from 1, 295 to 169, 998. In mock communities constructed from DNA input, significant positive relationships between the number of observations and sequencing depth (reads retained per sample) were observed in the 18S and shotgun datasets (**Figure 4**, green lines; both *p* <0.05). In contrast, such relationship was weaker and not statistically significant in the 28S dataset (*p* = 0.08). For mock communities constructed from individual specimens, the number of ASVs recovered per sample was significantly related to sequencing depth in both the 18S and 28S metabarcoding datasets (**Figure 4**, purple lines; both *p* <0.05), whereas no significant relationship was evident in the shotgun dataset. Additionally, the number of ASVs or contigs recovered varied among the 8 target nematode genera (**Supplementary Figure S2**).

**Figure 4.**
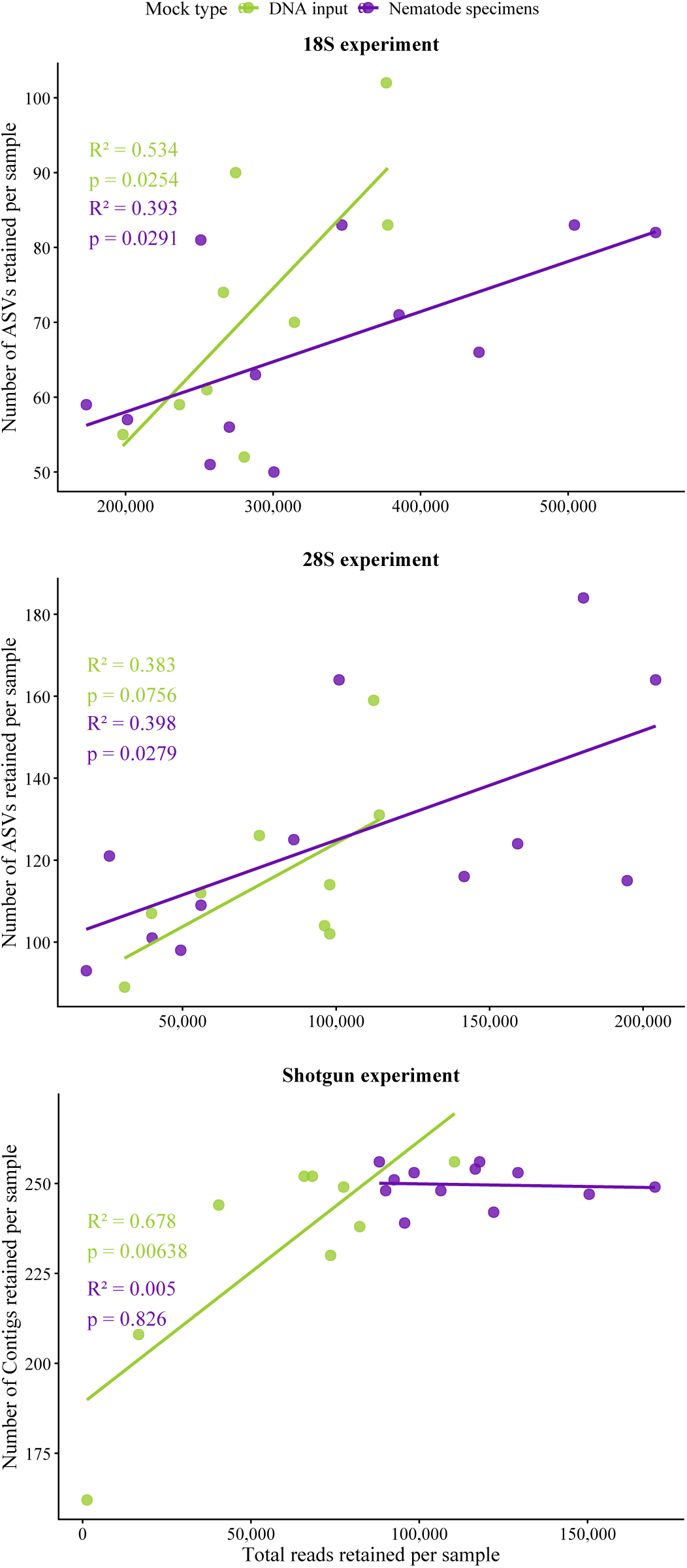
Relationship between number of observations (total number of ASVs or contigs) and sequencing depth (reads retained per sample). The coefficient of determination (R^2^) and *p*-value from linear regression are shown in the figure. The number of ASVs recovered across target nematodes is presented in **Supplementary Figure S2**.

For the detection of false negatives, two genera were not detected in some mock communities in which they were expected to occur. Across the 21 mock communities, *Leptolaimus* was not detected in 16 communities in the 28S dataset or in 2 communities in the 18S dataset, but was detected in all shotgun samples. *Paracanthonchus* was not detected in 2 communities in the 28S dataset. Among the 453 18S ASVs assigned to Nematoda, 416 were affiliated with the target nematode taxa, while the remaining 37 were classified as false positives (**Supplementary Table S3 and Figure S3**). The 28S dataset comprised 630 ASVs assigned to Nematoda, all of which were assigned to the target nematodes (**Supplementary Table S4**). In the shotgun dataset, 49 contigs out of 262 contigs were identified as false positives (**Supplementary Table S5**). The fraction of total non-target reads (including both “Other nematodes” and “Non nematodes”) was significantly larger in the shotgun dataset than in the 18S dataset and the 28S dataset (Wilcoxon test, *P* < 0.001; **Supplementary Figure S3**).

### Primer selection influences the consistency of beta diversity

The beta-diversity (between-sample comparison) of the mock communities derived from the same input material was similar across sequencing methods (**Supplementary Figure S4**) (Mantel test, all *P* <0.05; **Supplementary Table S6**). However, when comparing the observed read abundances relative to the expected proportions in both types of mock communities, shotgun sequencing and the 18S metabarcoding consistently yielded the lower pairwise Bray-Curtis dissimilarity than the 28S dataset. For the shotgun the dissimilarities ranged from 0.28 to 0.42 in the DNA input communities, and from 0.38 to 0.52 in the nematode specimens-composed communities and for 18S metabarcoding the range was 0.25 to 0.38 and 0.38 - 0. 59, respectively, while for 28S the range was 0.51 to 0.82 and 0.49 to 0.84 respectively (**Figure 5**). Despite these differences in dissimilarity values across sequencing methods, the deviation between expected proportions and observed abundances from the shotgun sequencing was statistically indistinguishable from that of 18S metabarcoding (**Figure 5**, Wilcox test, *P* > 0.05). In contrast, those deviations observed with 28S metabarcoding were significantly different from those obtained using either 18S metabarcoding or shotgun metagenomics in both types of mock communities (DNA input: *P* <0.01; nematode specimens: *P* <0.001). This finding suggests that 28S metabarcoding is less accurate in reflecting the expected community composition compared to 18S metabarcoding and shotgun.

**Figure 5.**
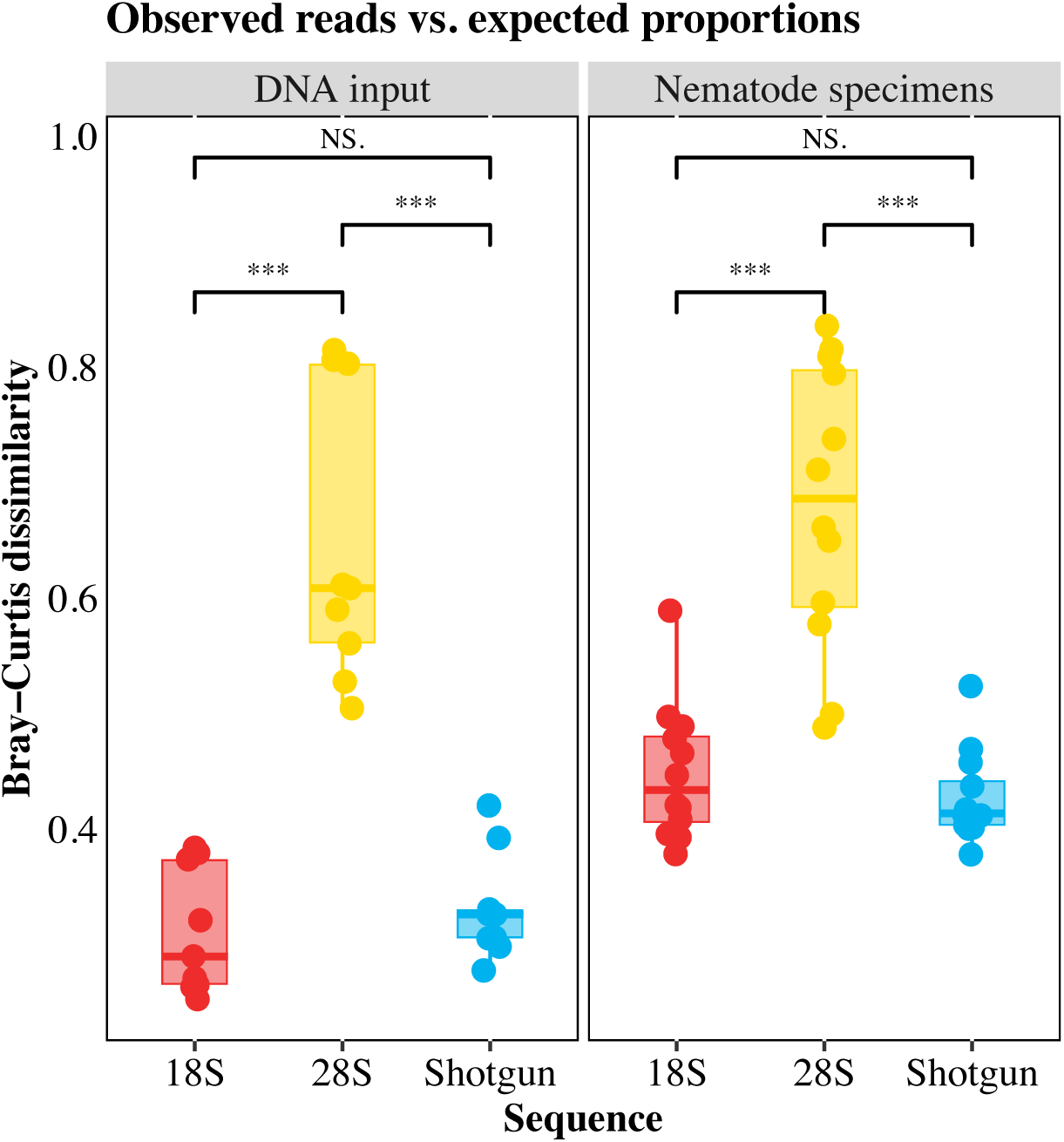
Bray-Curtis dissimilarity between mock community composition estimated from observed read abundances and the expected proportions across sequencing methods. Bray-Curtis dissimilarities were calculated at the genus-level. Significant differences in Bray-Curtis dissimilarity for each pairwise comparison were determined using a Wilcox test: ***, *P* <0.001; **, *P* <0.01; *, *P* <0.05; NS, no significance.

### 18S metabarcoding and shotgun sequencing reflect DNA quantity well

The composition of the three ecological groups, established based on their sensitivity to hypoxia, was compared in terms of relative abundances vs. the proportional amount of DNA in the mock communities as determined using the three sequencing methods (**Figure 6**). Overall, the abundance patterns of the three biological replicates (Tole_1, Tole_2, and Tole_3) for each community were very similar across the three sequencing methods (**Figure 6A**). For both the 18S and the shotgun datasets, the relative abundances of the ecological groups within each community well reflected the respective proportion of input DNA. In the case of 28S metabarcoding, however, the relative abundances of tolerant taxa were under-represented in all three levels of the mock communities and were not consistent with the proportions based on the quantity of input DNA (**Figure 6A**). A genus-level comparison showed that the observed relative abundances in the 18S dataset generally corresponded well with the expected proportions based on DNA input for *Sabatieria*, *Daptonema*, *Axonolaimus*, and *Desmolaimus* (**Figure 6B**). However, *Halomonhystera* was consistently overrepresented across the mock communities relative to its expected contribution, whereas *Leptolaimus*, *Paracanthonchus*, and *Microlaimus* were consistently underrepresented. In the 28S dataset, several genera, including *Sabatieria*, *Halomonhystera*, *Leptolaimus*, *Paracanthonchus*, and *Axonolaimus*, showed negligible relative abundances across all mock communities (**Figure 6B**), indicating poor recovery at this marker. In the shotgun dataset, *Paracanthonchus* was recovered in proportions that broadly reflected its expected DNA input. By contrast, *Halomonhystera*, *Leptolaimus*, and *Microlaimus* were underrepresented. Moreover, *Desmolaimus* also showed low relative read abundance, even in the sensitive-taxa-dominated communities where its DNA input exceeded that of the other nematode genera.

**Figure 6.**
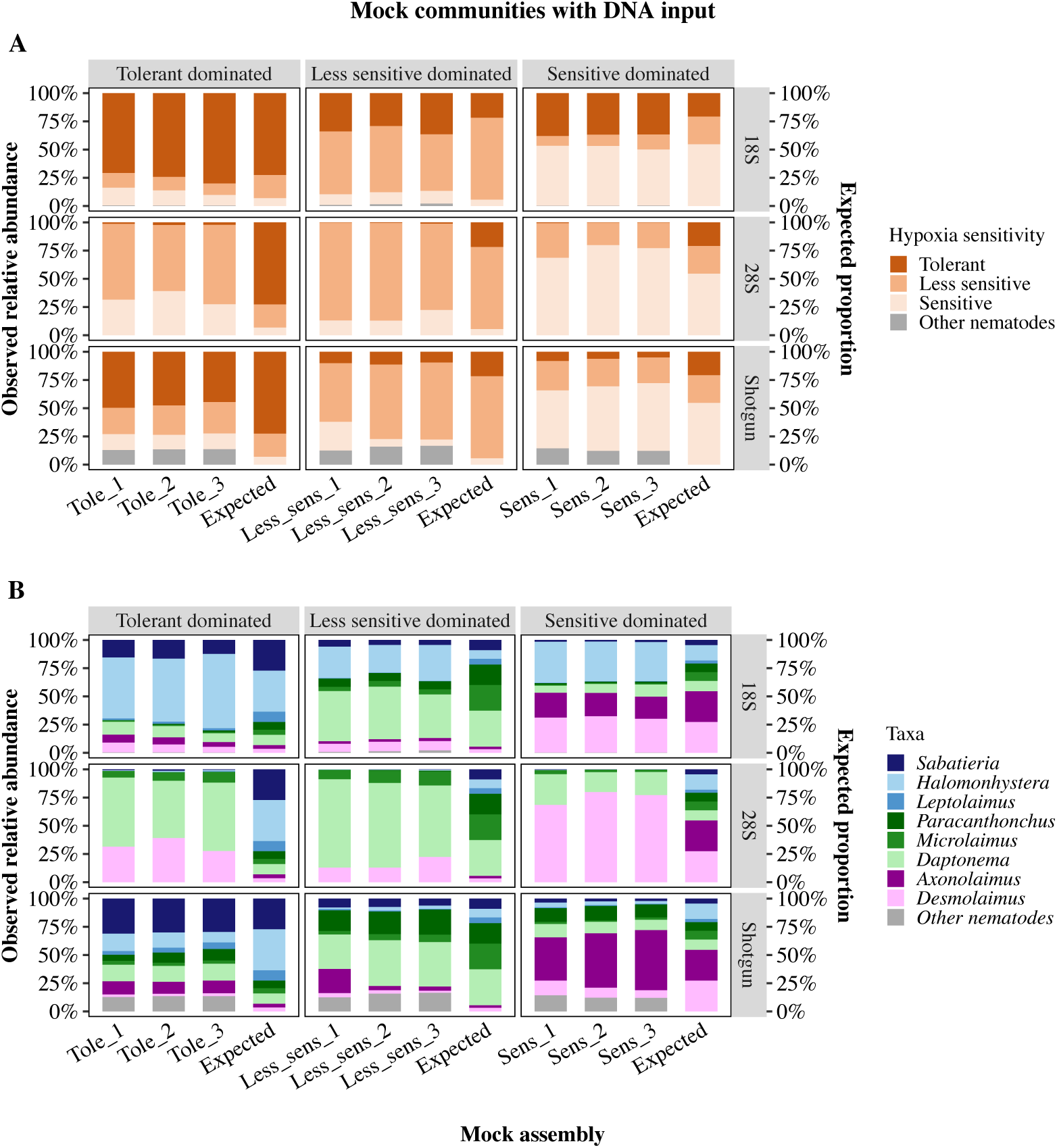
Abundance comparison of the mock community composed of inputted DNA; the observed relative abundance determined from sequences is shown on the left y-axis and the expected proportion of the total DNA quantity on the right y-axis. **A**) Proportional abundance in each mock community type (i.e., communities dominated by tolerant, less sensitive, and sensitive species) constructed from ecological groups of nematodes varying in their response to hypoxia. **B**) Taxonomic composition of the selected Nematoda genera in each mock community type. For both panels, Tole_1 to Tole_3, Less_sens_1 to Less_sens_3, and Sens_1, Sens_3 on the *x*-axis represent the biological triplicates of each mock community type, with “Tol”, “Les”, and “Sen” representing the expected composition of tolerant, less-sensitive, and sensitive communities based on the proportion of inputted DNA. In each panel, “Other nematodes” refers to ASVs in the 18S dataset or contigs in the shotgun dataset that could not be assigned to any of the target nematode genera, whereas “Non nematodes” refers to as ASVs assigned to other eukaryotes.

### Shotgun outperforms metabarcoding in quantifying the abundance relative to individual counts

Only in the shotgun dataset did the relative abundances of the three ecological groups correspond to the proportions of specimens added into the mock communities (**Figure 7A**). In both the 18S and 28S metabarcoding datasets, the less-sensitive group was underrepresented in the even mock communities. In addition, the tolerant-taxa group was consistently underrepresented across all mock communities in the 28S dataset.

**Figure 7.**
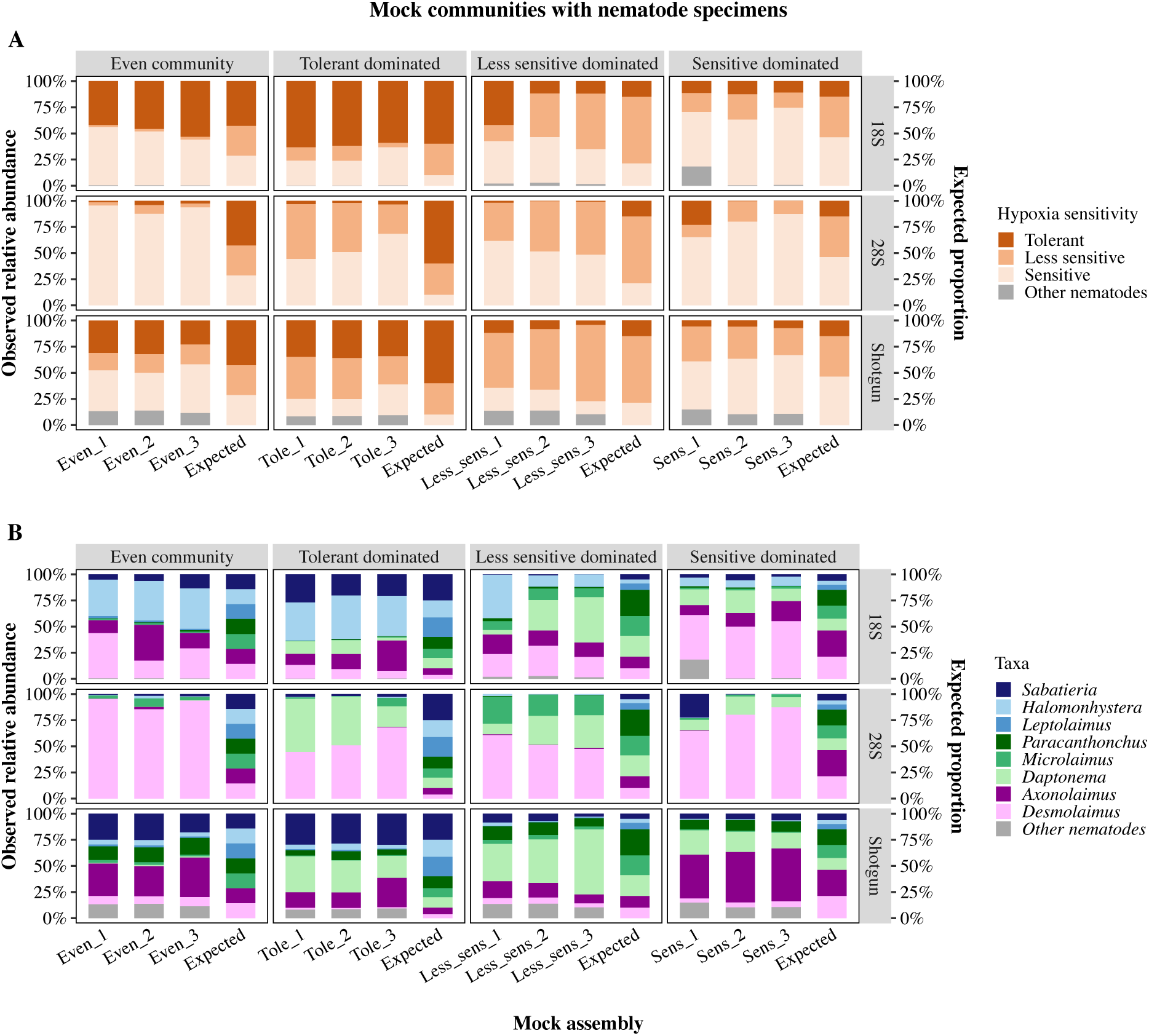
Abundance comparison for mock communities composed of individual nematode specimens: the observed relative abundance determined from sequences (left y-axis) vs. the expected proportion of total nematodes (right y-axis). **A**) Proportional abundance of ecological groups of nematodes varying in their response to hypoxia in each mock community (i.e., communities dominated by tolerant, less sensitive, and sensitive species). **B**) Taxonomic composition of the selected Nematoda genera in each mock community type. For both panels, Even_1 to Even_3, Tole_1 to Tole_3, Less_sens_1 to Less_sens_3, and Sens_1-Sens_3 on the x-axis represent the biological triplicates of each mock community type; “Even”, “Tol”, “Les”, and “Sen” represent the community composition expected from the proportion of inputted DNA. In each panel, “Other nematodes” refers to ASVs in the 18S dataset or contigs in the shotgun dataset that could not be assigned to any of the target nematode genera

Nematode composition within the even communities was skewed towards high abundances of *Halomonhystera* and *Desmolaimus* according to the 18S and 28S datasets, respectively; the estimated biomass of both genera was high (**Figure 7B**). However, according to the shotgun dataset the relative abundances of most of the genera (5 out of 8) were similar, as expected from the equal specimen numbers. Unexpectedly, *Daptonema* was also detected in the even communities by three methods, where it was not part of the input composition (**Figure 7B**). However, its relative abundance was very low throughout, suggesting that general laboratory contamination is unlikely and this detection likely resulted from low-level cross-sample carryover during sequencing.

Comparisons of nematode genus-level relative abundances within the staggered mock communities showed that the expected distributions based on the DNA input and specimen counts were broadly similar for most of the component genera (**Figures 6B** vs. **7B**). For instance, *Sabatieria* and *Halomonhystera* were dominated in the “Tolerant dominated” communities, where they were represented by higher numbers of specimens, and this pattern was largely reflected in the 18S dataset. In the 28S dataset, *Desmolaimus* was overrepresented relative to its expected proportion based on individual counts. In the shotgun dataset *Halomonhystera* was consistently underrepresented. Overall, *Leptolaimus* and *Microlaimus* showed negligible read abundances across all mock communities and sequencing methods, with the most pronounced underrepresentation observed in the shotgun dataset.

### 18S metabarcoding and shotgun well predict genus-specific abundance across samples

The correlations between the observed relative abundances vs. the expected proportion for a given genus illustrate how well the observed read abundance track the expected DNA input or specimen counts (**Figure 8**). In other words, whether the method can recover relative difference in genus abundance across samples. Those correlations differed depending on the sequencing approach and nematode genus. For the mock communities comprising DNA input, all target nematode genera exhibited significantly positive correlations between relative abundance and expected proportion, with the strongest relationships observed in the 18S and shotgun datasets (**Figure 8A**, all *P* <0.001) and weaker correlations in the 28S dataset (**Supplementary Table S7; Supplementary Figure S5A–C**). Overall, these results suggested that changes in DNA quantity, whether in the mock communities or in the library preparation pool and thus prior to sequencing, impact the abundance of each taxon and therefore its representation in a given community.

**Figure 8.**
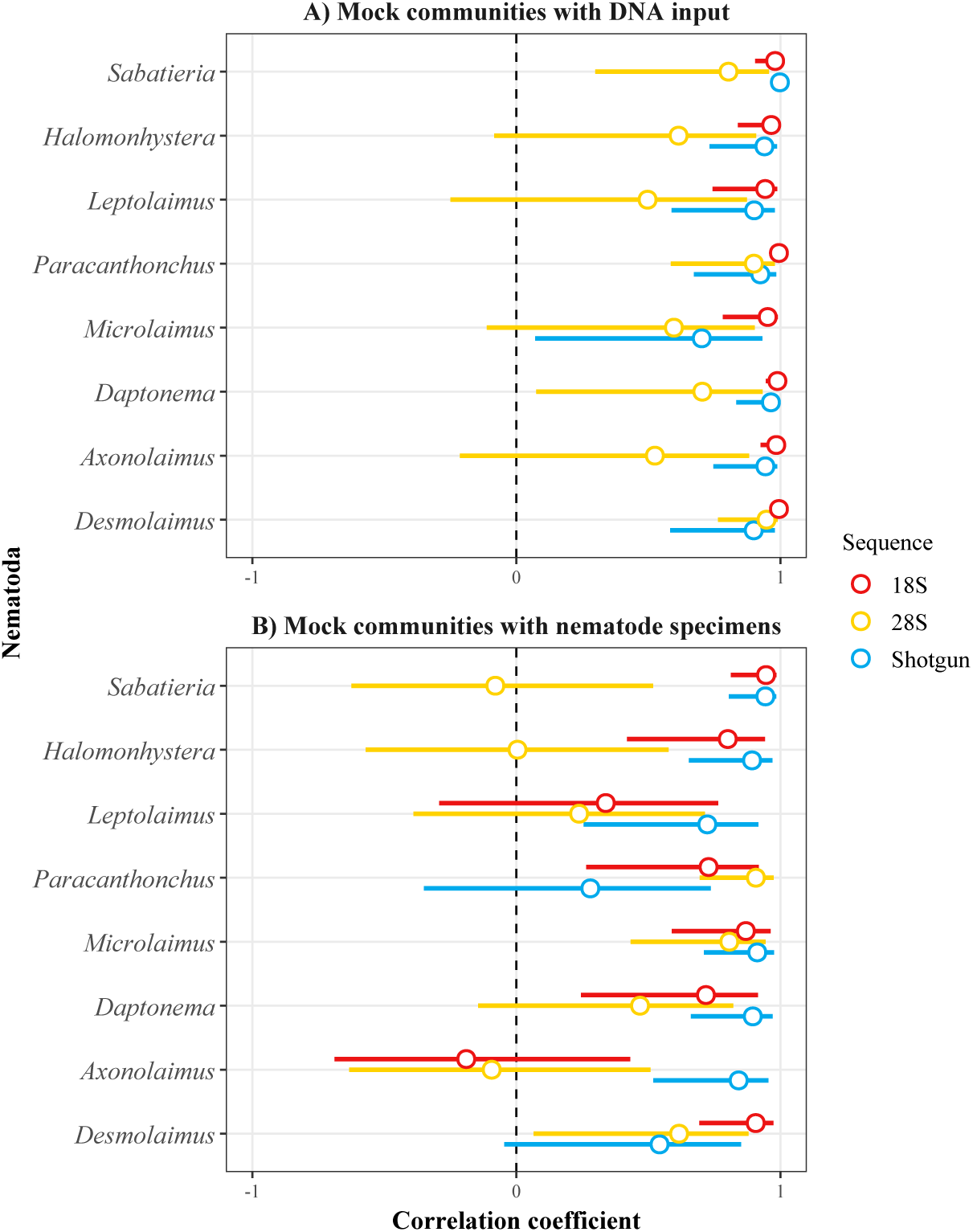
Coefficient plots of the Pearson correlation explaining the relationships between the relative abundances of each of the eight genera as measured from sequences and their expected proportion in the mock communities as determined by 18S metabarcoding (18S, red), 28S metabarcoding (28S, yellow), and shotgun sequencing (Shotgun, blue). **A**) Coefficients of the observed vs. expected values for the mock communities composed of DNA input. **B**) Coefficients of the observed vs. expected values for the mock communities composed of nematode individuals. The coefficients and their intervals are reported as one and two standard errors of the coefficient. A positive coefficient (> 0) indicates a positive correlation of the relative abundance of nematode genera with the expected proportion of either their specimen number or their DNA quantity in the mock communities. Conversely, a negative coefficient (< 0) indicates a negative correlation. A correlation coefficient near 1 or −1 indicates a strong linear relationship, and a correlation coefficient near 0, a weak linear relationship. The coefficients and *P*-values of each correlation for each genus are reported in Supplementary Figure S3 A–F, and the 95% confidence intervals for Pearson correlations coefficients in Supplementary **Table S7**.

Significant positive correlations between the observed relative abundances and the expected proportions based on the specimen counts were found for most genera in the nematode mock communities. Specifically, 6 out of 8 genera showed significant correlations with 18S metabarcoding (except *Leptolaimus* and *Axonolaimus*), while 7 genera were recovered using shotgun sequencing (except *Paracanthonchus* and *Desmolaimus* with *P* >0.05) (**Figure 8B**; **Supplementary Table S7**). In the 28S dataset, only the observed relative abundances of *Paracanthonchus*, *Microlaimus*, and *Desmolaimus* correlated with the expected proportion in terms of specimen number (*P* <0.05; Supplementary **Table S7**; **Supplementary Figure S5 E**). The strength of the correlations between the two types of mock communities showed stronger linear relationships in the mock communities comprising DNA input than nematode specimens, evidenced in the former by correlation coefficients near 1 (**Figure 8**).

## Discussion

High-throughput sequencing has advanced both biodiversity discovery and ecological monitoring, but its accuracy in quantifying species abundance remains uncertain and subject to ongoing debate (Bista et al., 2018; Latz et al., 2022; Schmidt et al., 2022). This study evaluated the accuracy and precision of DNA metabarcoding and shotgun sequencing for quantifying nematode abundances. Although DNA extraction was required in both experiments to assess the quantitative performance of the sequencing approaches, we also evaluated whether this performance was influenced by differences in the starting material used to construct the mock communities: DNA quantities in Experiment 1 and known specimen counts in Experiment 2.

In nematodes, part of the intraspecific variation in rRNA gene is attributed to intragenomic variation (Bik et al., 2013; Qing et al., 2020), whereby a single nematode individual contains multiple, slightly different copies of the same rRNA gene within its genome. However, the extent of variation in 18S rRNA or 28S rRNA genes within and among nematode genus or species, as well as the relative abundance of individual variants, remains poorly understood. In this study, the strength of the positive relationships between observed richness (number of ASVs / contigs retained) and sequencing depth varied across primer choice and mock community types, and multiple ASVs / contigs were recovered for each target genus. These findings suggest that intragenomic rRNA variation contributes to observed diversity patterns, with increased sequencing depth enhancing the detection of these variants. This interpretation is consistent with previous assumption that rRNA gene copy number is not constant within nematode species and may vary under evolutionary pressure (Bik et al., 2013; Qing et al., 2020). As a result, a single nematode species can harbor multiple non-identical copies of the 18S and 28S rRNA genes and therefore give rise to several distinct ASVs. For example, < 100 nematode ASVs have been reported per individual nematode in 18S datasets processed by DADA2 (Pereira et al., 2020), and similar patterns have been observed in earlier studies (Porazinska et al., 2009, 2010).

Previous work demonstrated that metabarcoding and shotgun data produce qualitatively similar community profiles of metazoans (Arribas et al., 2016; Bista et al., 2018; Ji et al., 2020). Our data supported hypothesis 1 and are consistent with studies reporting the comparable β-diversity obtained by metabarcoding and shotgun metagenomics, with both able to recover signals indicative of community structural changes. In both experiments, 18S metabarcoding and shotgun sequencing produced more similar mock community composition than 28S metabarcoding, likely due to differences in primer coverage. The 18S rRNA gene is a more effective marker than the 28S rRNA gene in the detection of the diversity of nematodes in soil (Ahmed et al., 2019; Floyd et al., 2002) and in marine systems (Avo et al., 2017; Pereira et al., 2010), likely because of its broader taxonomic coverage and more extensive representation in reference databases (Bik et al., 2013; Charrier et al., 2024). Additionally, although 18S and 28S rRNA genes are not expected to differ substantially in copy number within the same genome, read abundances assigned to the same taxon may vary between 18S and 28S metabarcoding libraries due to marker-specific amplification biases, primer mismatches, and differences in reference database coverage or taxonomic assignment accuracy. Such biases can affect the estimated abundances of dominant taxa that could be interpreted as community-level dissimilarities.

A determination of the false-negative and false-positive detection rates in high-throughput sequencing data is critical for accurate taxon identification (Bell et al., 2021; Bista et al., 2018; Schmidt et al., 2022; Sun et al., 2023). Studies of invertebrate communities have shown that metabarcoding and shotgun sequencing produce comparable false-negative detection rates, while shotgun sequencing is prone to a high rate of false-positive detections (Bista et al., 2018; Schmidt et al., 2022). All three sequencing methods detected the target nematodes in most mock communities; however, false-negative detection rate varied slightly among methods, with the rate observed for *Leptolaimus* in the 28S dataset. However, compared with the 18S metabarcoding, the shotgun approach generated significantly more false positives (i.e., sequence reads assigned to Other nematodes), despite both approaches being applied to the same mock samples. This implies that the false-positives detected in the shotgun dataset were unlikely to result from web-lab contamination (e.g., from reagents or sample handling), but were instead more likely caused by computational misclassification and traces of other nematodes already present within or associated with the target nematodes. For example, several false-positives in the shotgun dataset were taxonomically annotated as the close relatives of *Axonolaimus*, including *Odontophora* and *Araeolaimus* (**Supplementary Table S5**). In addition, other false-positive contigs were assigned to nematodes that may represent ingested material associated with the target taxa and are vertebrate parasites. A possible explanation is that DNA from these taxa was introduced into the sediment through human and agricultural waste inputs at the sampling sites, and was subsequently detected indirectly through ingested material associated with the target nematodes. Nevertheless, most of these false-positives accounted for less than 0.5 % of the total reads within individual communities, which falls within the acceptable detection range reported by Ye et al. (2019). These authors also emphasized that, when applying shotgun metagenomics to environmental samples, low-abundance taxonomic assignments should be interpreted with caution or filtered out using stringent post-classification thresholds. Alternatively, taxonomic classification can be cross-validated using multiple reference databases.

Support for hypothesis 2 was conditional on the primer choice and the DNA input. 28S metabarcoding was skewed towards high abundances of *Daptonema* and *Desmolaimus* in mock communities constructed using DNA input, suggesting primer-driven overestimation of these genera. Shotgun sequencing outperformed 18S metabarcoding in estimating the abundance of large-bodied organisms with estimated high gene copy numbers. Despite similar DNA inputs of *Halomonhystera* and *Leptolaimus*, differences in the 18S rRNA gene copy numbers per genome, i.e., higher in *Halomonhystera* than in *Leptolaimus* (Fonseca and Decraemer, 2008; Holovachov and Boström, 2013), likely explain their contrasting relative abundances. PCR-based metabarcoding may further amplify these differences, distorting the relationship between sequence reads and taxon abundance (Martin et al., 2022). Accordingly, the overestimation of *Halomonhystera* relative to its DNA input highlights the risks of overinterpreting read abundance as taxon abundance in metabarcoding datasets, particularly for taxa with high rRNA gene copy numbers; however, this bias was not evident in the shotgun dataset. Notably, the 18S rRNA reads were also extracted and reconstructed from shotgun data in our study, with contigs assembled from these sequences rather than PCR-amplified fragments. This approach reduces the influence of marker-specific biases and PCR amplification bias. Therefore, shotgun sequencing provided a more consistent representation of the relationship between observed relative reads and input DNA compared to metabarcoding.

Hypothesis 3 was partially supported by Experiment 2, where the shotgun data corresponded well with the number of individuals only for taxa of moderate to large body size. In the even mock communities, *Halomonhystera* and *Desmolaimus* were consistently over-represented in the 18S and 28S datasets, respectively, mirroring patterns observed in Experiment 1. Their larger body size likely contributes to higher biomass (Górska et al., 2020; Górska and Wlodarska-Kowalczuk, 2017). This, together with potential differences in rRNA gene copy number, likely explains their preferential amplification. In contrast, the underrepresentation of *Leptolaimus*, *Microlaimus*, and *Paracanthoncus* in both metabarcoding datasets was mostly likely due to primer mismatches. Additionally, the relative thick cuticle of *Paracanthoncus* may hinder lysis and DNA extraction, further reducing amplification efficiency. Thus, even the number of nematode individuals were equal in the mock communities, those contributing more to the overall biomass may be more likely to dominate amplicon read counts, and effect that appeared more pronounced for 18S than for 28S (**Supplementary Table S6**). Although body size is one of the most commonly used proxies for biomass (Peters, 1983; Saint-Germain et al., 2007), other biological traits may also influence biomass estimates. Variation in gonad cell number, reproductive strategies, and development stage may affect the number of rDNA copies contributed by an individual specimen, thereby leading to differences in relative read abundances among species. For example, *Halomonhystera* is ovoviviparous and may therefore contain considerably more nuclei than other species of comparable size. Its relatively large gonads may further increase rDNA copy number, potentially contributing to its high read abundance. Nonetheless, read abundance recovered from the shotgun data was less biased with respect to estimated body size, particularly for larger-bodied taxa.

Both metabarcoding and shotgun sequencing proved challenging in quantifying the abundance of *Leptolaimus* and *Microlaimus*, two genera with a small body size. This highlights the importance of sufficient sequencing depth to improve the detection of small-bodied and low-abundance taxa, which contribute limited amounts of DNA to bulk samples and may otherwise be underrepresented in read data. In our study, density extraction of benthic meiofauna offered a more concentrated community sample and, thus, a higher DNA yield than direct extraction from sediment. Metabarcoding does not require high-quality genetic material for PCR amplification, whereas DNA of high quality and quantity is a prerequisite for successful library preparation in shotgun sequencing (Wilcox et al., 2018). The latter method is therefore limited when used to quantify organisms present at low DNA concentrations and may not be suitable for eDNA surveillance, such as applied in the detection and monitoring of rare or threatened organisms in aquatic systems (Zhan and MacIsaac, 2015). Moreover, processing shotgun sequences requires comprehensive genome references, which may limit its suitability for long-term environmental biomonitoring.

Several lines of evidence from our study indicate that input material influenced quantitative performance, as demonstrated by the correlation analyses. Relative read data from both metabarcoding and shotgun sequencing generally tracked DNA quantity more closely than specimen counts, suggesting better quantitative recovery in Experiment 1, where mock communities were constructed using defined DNA input. This discrepancy was likely due to variation in nematode body size among genera. That means that specimen counts may not reliably reflect biomass or DNA contribution when target organisms vary significantly in size, as previously noted (Elbrecht and Leese, 2015). Therefore, DNA quantity may represent a more suitable a more suitable input material for future mock community studies involving organisms with variable body sizes, as it better predicts sequencing read abundances.

## Conclusions

Our study shows that, overall, 18S metabarcoding and shotgun sequencing are comparably effective in detecting marine nematodes and in reflecting the overall community structure across sampling sites. Specifically, their quantitative performance was similar for within-taxon abundance and in occurrence evaluations, whereases shotgun sequencing proved more reliable for across-taxon quantification within samples. If the aim of the research is to determine which organisms are present in environmental samples, metabarcoding is a cost-effective method with a low rate of false positives. The 28S primers used in the experiments, however, introduce greater bias and variability in estimating metabarcoding read abundances, making it a less reliable primer set than 18S for quantitative community profiling of marine nematodes. Both mock community experiments clearly demonstrate that shotgun sequencing provides more consistent quantitative performance than metabarcoding, especially when assessing samples containing individuals with large size differences. While shotgun sequencing holds promise for quantifying eukaryotic communities, its practical application is currently constrained by the limited representation of eukaryotic genomes in reference databases, particularly for marine nematodes and other meiofauna.

In summary, the genus-level biases detected in our study have important implications for monitoring, as benthic ecological status assessments often depend on accurate estimates of the relative abundances of nematode ecological groups. Distortion in read abundance relative to expected proportion indicated an overrepresentation of hypoxia tolerant nematodes, which may falsely signal a healthy habitat, and delays timely responses to environmental degradation (e.g., declining sediment quality). Conversely, the underestimation of sensitive taxa obscures early signals of ecosystem stress, potentially leading to misclassification of ecological status and misdirected management interventions. Thus, to ensure the quantitative reliability of nematode metabarcoding in environmental monitoring, it is crucial to identify and address biases through the calibration with mock community and real samples, the use of standardized protocols, and the continual improvement of reference databases. Once these measures are in place, the studied nematode ecological groups could serve as model organisms, supporting the application of metabarcoding for robust benthic ecological status assessment and long-term environmental monitoring.

## Acknowledgements

The authors thank Ola Svensson, Caroline Raymond, and Jonas Gunnarsson for assistance during sampling, and Johanna Hehberg for support during the experiment setup. We also acknowledge support from the National Genomics Infrastructure in Stockholm funded by Science for Life Laboratory, the Knut and Alice Wallenberg Foundation and the Swedish Research Council, and SNIC/Uppsala Multidisciplinary Center for Advanced Computational Science for assistance with massively parallel sequencing and access to the UPPMAX computational infrastructure.

## Author contributions

Dandan Izabel-Shen conceived the study, designed and conducted the experiments, processed the samples, performed the bioinformatic analyses, analyzed the data, and wrote the manuscript. Henrik Sandberg contributed to the manuscript writing, bioinformatic analyses, data analyses. Mohammed Ahmed assisted with experimental design and the morphological identification of the nematodes. Elias Broman performed the bioinformatic analyses of the shotgun sequencing data. Oleksandr Holovachov contributed to the experimental design and conducted the morphological identification of the nematodes. Francisco J. A. Nascimento conceived and financed the study, designed the experiments, and contributed to manuscript writing. All authors discussed the results and commented on the manuscript.

## Data availability

The FASTQ files and associated metadata have been made available in the European Nucleotide Archive under the accession number PRJEB98572.

## Code availability

Our R scripts for statistics, data visualization and computing notes are available in GitHub (https://github.com/IzabelShen/Evaluating-quantitative-ability-of-metabarcoding-vs.-shotgun-in-marine-nematodes).

## References

Ahmed, M., Back, M.A., Prior, T., Karssen, G., Lawson, R., Adams, I., Sapp, M., 2019. Metabarcoding of soil nematodes: the importance of taxonomic coverage and availability of reference sequences in choosing suitable marker(s). Metabarcoding and Metagenomics 3, e36408. 10.3897/mbmg.3.36408

Andrews, S., 2010. FastQC: A quality control tool for high throughput sequence data.

Arribas, P., Andújar, C., Hopkins, K., Shepherd, M., Vogler, A.P., 2016. Metabarcoding and mitochondrial metagenomics of endogean arthropods to unveil the mesofauna of the soil. Methods in Ecology and Evolution 7, 1071–1081. 10.1111/2041-210X.12557

Avo, A., Daniell, T., Neilson, R., Oliveira, S., Branco, J., Adao, H., 2017. DNA Barcoding and Morphological Identification of Benthic Nematodes Assemblages of Estuarine Intertidal Sediments: Advances in Molecular Tools for Biodiversity Assessment. Front. Mar. Sci. 4. 10.3389/fmars.2017.00066

Bell, K.L., Petit III, R.A., Cutler, A., Dobbs, E.K., Macpherson, J.M., Read, T.D., Burgess, K.S., Brosi, B.J., 2021. Comparing whole-genome shotgun sequencing and DNA metabarcoding approaches for species identification and quantification of pollen species mixtures. Ecology and Evolution 11, 16082–16098. 10.1002/ece3.8281

Bik, H.M., Fournier, D., Sung, W., Bergeron, R.D., Thomas, W.K., 2013. Intra-Genomic Variation in the Ribosomal Repeats of Nematodes. PLOS ONE 8, e78230. 10.1371/journal.pone.0078230

Bik, H.M., Porazinska, D.L., Creer, S., Caporaso, J.G., Knight, R., Thomas, W.K., 2012. Sequencing our way towards understanding global eukaryotic biodiversity. Trends in Ecology & Evolution 27, 233–243. 10.1016/j.tree.2011.11.010

Bista, I., Carvalho, G.R., Tang, M., Walsh, K., Zhou, X., Hajibabaei, M., Shokralla, S., Seymour, M., Bradley, D., Liu, S., Christmas, M., Creer, S., 2018. Performance of amplicon and shotgun sequencing for accurate biomass estimation in invertebrate community samples. Mol Ecol Resour 18, 1020–1034. 10.1111/1755-0998.12888

Bolger, A.M., Lohse, M., Usadel, B., 2014. Trimmomatic: A flexible trimmer for Illumina sequence data. Bioinformatics 30, 2114–2120. 10.1093/bioinformatics/btu170

Bonaglia, S., Nascimento, F.J.A., Bartoli, M., Klawonn, I., Brüchert, V., 2014. Meiofauna increases bacterial denitrification in marine sediments. Nat Commun 5, 5133. 10.1038/ncomms6133

Broman, E., Izabel-Shen, D., Rodriguez-Gijon, A., Bonaglia, S., Garcia, S., Nascimento, F., 2022. Microbial functional genes are driven by gradients in sediment stoichiometry, oxygen, and salinity across the Baltic benthic ecosystem. Microbiome 10, 126. 10.1186/s40168-022-01321-z

Broman, E., Raymond, C., Sommer, C., Gunnarsson, J.S., Creer, S., Nascimento, F.J.A., 2019. Salinity drives meiofaunal community structure dynamics across the Baltic ecosystem. Molecular Ecology 28, 3813–3829. 10.1111/mec.15179

Callens, M., Le Berre, G., Van den Bulcke, L., Lolivier, M., Derycke, S., 2025. An Accessible Metagenomic Strategy Allows for Better Characterisation of Invertebrate Bulk Samples. Molecular Ecology Resources 25, e14126. 10.1111/1755-0998.14126

Charrier, E., Chen, R., Thundathil, N., Gilleard, J., 2024. A set of nematode rRNA cistron databases and a primer assessment tool to enable more flexible and comprehensive metabarcoding. Molecular Ecology Resources 24, e13965. 10.1111/1755-0998.13965

Conley, D.J., Carstensen, J., Aigars, J., Axe, P., Bonsdorff, E., Eremina, T., Haahti, B.-M., Humborg, C., Jonsson, P., Kotta, J., Lännegren, C., Larsson, U., Maximov, A., Medina, M.R., Lysiak-Pastuszak, E., Remeikaitė-Nikienė, N., Walve, J., Wilhelms, S., Zillén, L., 2011. Hypoxia Is Increasing in the Coastal Zone of the Baltic Sea. Environ. Sci. Technol. 45, 6777–6783. 10.1021/es201212r

Cook, L.S.J., Briscoe, A.G., Fonseca, V.G., Boenigk, J., Woodward, G., Bass, D., 2025. Microbial, holobiont, and Tree of Life eDNA/eRNA for enhanced ecological assessment. Trends in Microbiology 33, 48–65. 10.1016/j.tim.2024.07.003

Creer, S., Fonseca, V.G., Porazinska, D.L., Giblin-Davis, R.M., Sung, W., Power, D.M., Packer, M., Carvalho, G.R., Blaxter, M.L., Lambshead, P.J.D., Thomas, W.K., 2010. Ultrasequencing of the meiofaunal biosphere: practice, pitfalls and promises. Molecular Ecology 19, 4–20. 10.1111/j.1365-294X.2009.04473.x

Deagle, B.E., Thomas, A.C., McInnes, J.C., Clarke, L.J., Vesterinen, E.J., Clare, E.L., Kartzinel, T.R., Eveson, J.P., 2019. Counting with DNA in metabarcoding studies: How should we convert sequence reads to dietary data? Mol Ecol 28, 391–406. 10.1111/mec.14734

di Montanara, A., Baldrighi, E., Franzo, A., Catani, L., Grassi, E., Sandulli, R., Semprucci, F., 2022. Free-living nematodes research: State of the art, prospects, and future directions. A bibliometric analysis approach. Ecological Informatics 72, 101891. 10.1016/j.ecoinf.2022.101891

Elbrecht, V., Leese, F., 2015. Can DNA-Based Ecosystem Assessments Quantify Species Abundance? Testing Primer Bias and Biomass—Sequence Relationships with an Innovative Metabarcoding Protocol. PLOS ONE 10, e0130324. 10.1371/journal.pone.0130324

Elliott, L., Coissac, E., 2025. Can Amplicon Sequencing Be Replaced by Metagenomics for Biodiversity Inventories? Molecular Ecology Resources 25, e70047. 10.1111/1755-0998.70047

Ershova, E., Wangensteen, O., Descoteaux, R., Barth-Jensen, C., Praebel, K., 2021. Metabarcoding as a quantitative tool for estimating biodiversity and relative biomass of marine zooplankton. ICES Journal of Marine Science 78, 3342–3355. 10.1093/icesjms/fsab171

Ewels, P., Magnusson, M., Lundin, S., Käller, M., 2016. MultiQC: summarize analysis results for multiple tools and samples in a single report. Bioinformatics 32, 3047–3048. 10.1093/bioinformatics/btw354

Floyd, R., Abebe, E., Papert, A., Blaxter, M., 2002. Molecular barcodes for soil nematode identification. Molecular Ecology 11, 839–850. 10.1046/j.1365-294X.2002.01485.x

Fonseca, G., Decraemer, W., 2008. State of the art of the free-living marine Monhysteridae (Nematoda). Journal of the Marine Biological Association of the United Kingdom 88, 1371–1390. 10.1017/S0025315408001719

Fonseca, V.G., Carvalho, G.R., Sung, W., Johnson, H.F., Power, D.M., Neill, S.P., Packer, M., Blaxter, M.L., Lambshead, P.J.D., Thomas, W.K., Creer, S., 2010. Second-generation environmental sequencing unmasks marine metazoan biodiversity. Nat Commun 1, 98. 10.1038/ncomms1095

Frontalini, F., Greco, M., Semprucci, F., Cermakova, K., Merzi, T., Pawlowski, J., 2025. Developing and testing a new Ecological Quality Status index based on marine nematode metabarcoding: A proof of concept. Chemosphere 370, 143992. 10.1016/j.chemosphere.2024.143992

Gielings, R., Fais, M., Fontaneto, D., Creer, S., Costa, F.O., Renema, W., Macher, J.-N., 2021. DNA Metabarcoding Methods for the Study of Marine Benthic Meiofauna: A Review. Front. Mar. Sci. 8, 730063. 10.3389/fmars.2021.730063

Górska, B., Soltwedel, T., Schewe, I., Włodarska-Kowalczuk, M., 2020. Bathymetric trends in biomass size spectra, carbon demand, and production of Arctic benthos (76- 5561 m, Fram Strait). Progress in Oceanography 186, 102370. 10.1016/j.pocean.2020.102370

Górska, B., Wlodarska-Kowalczuk, M., 2017. Food and disturbance effects on Arctic benthic biomass and production size spectra. Progress in Oceanography 152, 50–61. 10.1016/j.pocean.2017.02.005

Holovachov, O., 2016. Metabarcoding of marine nematodes - evaluation of reference datasets used in tree-based taxonomy assignment approach. BDJ 4, e10021. 10.3897/BDJ.4.e10021

Holovachov, O., Boström, S., 2013. Swedish Plectida (Nematoda). Part 4. The genus Leptolaimus de Man, 1876 Zootaxa 3739, 1–99. 10.11646/zootaxa.3739.1.1

Holovachov, O., Haenel, Q., Bourlat, S.J., Jondelius, U., 2017. Taxonomy assignment approach determines the efficiency of identification of OTUs in marine nematodes. R. Soc. open sci. 4, 170315. 10.1098/rsos.170315

Iburg, S., Izabel-Shen, D., Austin, A., Hansen, J., Eklof, J., Nascimento, F., 2021. Effects of Recreational Boating on Microbial and Meiofauna Diversity in Coastal Shallow Ecosystems of the Baltic Sea. mSphere 6, e00127–21. 10.1128/mSphere.00127-21

Ji, Y., Huotari, T., Roslin, T., Schmidt, N.M., Wang, J., Yu, D.W., Ovaskainen, O., 2020. SPIKEPIPE: A metagenomic pipeline for the accurate quantification of eukaryotic species occurrences and intraspecific abundance change using DNA barcodes or mitogenomes. Molecular Ecology Resources 20, 256–267. 10.1111/1755-0998.13057

Kennedy, A., Jacoby, C., 1999. Biological indicators of marine environmental health: Meiofauna - A neglected benthic component? Environ Monit Assess 54, 47–68. 10.1023/A:1005854731889

Kiu, R., 2017. blastoutput2gff. GitHub repository.

Kopylova, E., Noé, L., Touzet, H., 2012. SortMeRNA: fast and accurate filtering of ribosomal RNAs in metatranscriptomic data. Bioinformatics 28, 3211–3217. 10.1093/bioinformatics/bts611

Kuliński, K., Rehder, G., Asmala, E., Bartosova, A., Carstensen, J., Gustafsson, B., Hall, P.O.J., Humborg, C., Jilbert, T., Jürgens, K., Meier, H.E.M., Müller-Karulis, B., Naumann, M., Olesen, J.E., Savchuk, O., Schramm, A., Slomp, C.P., Sofiev, M., Sobek, A., Szymczycha, B., Undeman, E., 2022. Biogeochemical functioning of the Baltic Sea. Earth Syst. Dynam. 13, 633–685. 10.5194/esd-13-633-2022

Lamb, P., Hunter, E., Pinnegar, J., Creer, S., Davies, R., Taylor, M., 2019. How quantitative is metabarcoding: A meta-analytical approach. Molecular Ecology 28, 420–430. 10.1111/mec.14920

Langmead, B., Salzberg, S.L., 2012. Fast gapped-read alignment with Bowtie 2. Nat Methods 9, 357–359. 10.1038/nmeth.1923

Latz, M., Grujcic, V., Brugel, S., Lycken, J., John, U., Karlson, B., Andersson, A, Andersson, AF, 2022. Short- and long-read metabarcoding of the eukaryotic rRNA operon: Evaluation of primers and comparison to shotgun metagenomics sequencing. Molecular Ecology Resources 22, 2304–2318. 10.1111/1755-0998.13623

Li, D., Liu, C.-M., Luo, R., Sadakane, K., Lam, T.-W., 2015. MEGAHIT: an ultra-fast single-node solution for large and complex metagenomics assembly via succinct de Bruijn graph. Bioinformatics 31, 1674–1676. 10.1093/bioinformatics/btv033

Liao, Y., Smyth, G.K., Shi, W., 2014. featureCounts: an efficient general purpose program for assigning sequence reads to genomic features. Bioinformatics 30, 923–930. 10.1093/bioinformatics/btt656

Markmann, M., Tautz, D., 2005. Reverse taxonomy: an approach towards determining the diversity of meiobenthic organisms based on ribosomal RNA signature sequences. Philosophical Transactions of the Royal Society B: Biological Sciences 360, 1917–1924. 10.1098/rstb.2005.1723

Martin, J., Santi, I., Pitta, P., John, U., Gypens, N., 2022. Towards quantitative metabarcoding of eukaryotic plankton: an approach to improve 18S rRNA gene copy number bias. Metabarcoding and Metagenomics 6, 245–259. 10.3897/mbmg.6.85794

Modig, H., Ólafsson, E., 1998. Responses of Baltic benthic invertebrates to hypoxic events. Journal of Experimental Marine Biology and Ecology 229, 133–148. 10.1016/S0022-0981(98)00043-4

Nascimento, F.J.A., Näslund, J., Elmgren, R., 2012. Meiofauna enhances organic matter mineralization in soft sediment ecosystems. Limnology & Oceanography 57, 338– 346. 10.4319/lo.2012.57.1.0338

Nasko, D.J., Koren, S., Phillippy, A.M., Treangen, T.J., 2018. RefSeq database growth influences the accuracy of k-mer-based lowest common ancestor species identification. Genome Biology 19, 165. 10.1186/s13059-018-1554-6

Paula, D.P., Barros, S.K.A., Pitta, R.M., Barreto, M.R., Togawa, R.C., Andow, D.A., 2022. Metabarcoding versus mapping unassembled shotgun reads for identification of prey consumed by arthropod epigeal predators. Gigascience 11, giac020. 10.1093/gigascience/giac020

Pereira, T., Fonseca, G., Mundo-Ocampo, M., Guilherme, B., Rocha-Olivares, A., 2010. Diversity of free-living marine nematodes (Enoplida) from Baja California assessed by integrative taxonomy. Marine Biology 157, 1665–1678. 10.1007/s00227-010-1439-z

Pereira, T.J., De Santiago, A., Schuelke, T., Hardy, S.M., Bik, H.M., 2020. The impact of intragenomic rRNA variation on metabarcoding-derived diversity estimates: A case study from marine nematodes. Environmental DNA 2, 519–534. 10.1002/edn3.77

Peters, R.H., 1983. The Ecological Implications of Body Size, Cambridge Studies in Ecology. Cambridge University Press, Cambridge. 10.1017/CBO9780511608551

Piñol, J., Mir, G., Gomez-Polo, P., Agustí, N., 2015. Universal and blocking primer mismatches limit the use of high-throughput DNA sequencing for the quantitative metabarcoding of arthropods. Molecular Ecology Resources 15, 819–830. 10.1111/1755-0998.12355

Pinto, R., Patrício, J., Baeta, A., Fath, B.D., Neto, J.M., Marques, J.C., 2009. Review and evaluation of estuarine biotic indices to assess benthic condition. Ecological Indicators 9, 1–25. 10.1016/j.ecolind.2008.01.005

Porazinska, D., Giblin-Davis, R., Faller, L., Farmerie, W., Kanzaki, N., Morris, K., Powers, T., Tucker, A., Sung, W., Thomas, W., 2009. Evaluating high-throughput sequencing as a method for metagenomic analysis of nematode diversity. MOLECULAR ECOLOGY RESOURCES 9, 1439–1450. 10.1111/j.1755-0998.2009.02611.x

Porazinska, D.L., Giblin-Davis, R.M., Esquivel, A., Powers, T.O., Sung, W., Thomas, W.K., 2010. Ecometagenetics confirm high tropical rainforest nematode diversity. Molecular Ecology 19, 5521–5530. 10.1111/j.1365-294X.2010.04891.x

Qing, X., Bik, H., Yergaliyev, T.M., Gu, J., Fonderie, P., Brown-Miyara, S., Szitenberg, A., Bert, W., 2020. Widespread prevalence but contrasting patterns of intragenomic rRNA polymorphisms in nematodes: Implications for phylogeny, species delimitation and life history inference. Molecular Ecology Resources 20, 318–332. 10.1111/1755-0998.13118

R Core Team, 2021. R: A language and environment for statistical computing.

Ridall, A., Ingels, J., 2021. Suitability of Free-Living Marine Nematodes as Bioindicators: Status and Future Considerations. Front. Mar. Sci. 8, 685327. 10.3389/fmars.2021.685327

Schenk, J., Fontaneto, D., 2020. Biodiversity analyses in freshwater meiofauna through DNA sequence data. Hydrobiologia 847, 2597–2611. 10.1007/s10750-019-04067-2

Schenk, J., Geisen, S., Kleinboelting, N., Traunspurger, W., 2019. Metabarcoding data allow for reliable biomass estimates in the most abundant animals on earth. Metabarcoding and Metagenomics 3, e46704. 10.3897/mbmg.3.46704

Schmidt, A., Schneider, C., Decker, P., Hohberg, K., Römbke, J., Lehmitz, R., Bálint, M., 2022. Shotgun metagenomics of soil invertebrate communities reflects taxonomy, biomass, and reference genome properties. Ecology and Evolution 12. 10.1002/ece3.8991

Schratzberger, M., Ingels, J., 2018. Meiofauna matters: The roles of meiofauna in benthic ecosystems. Journal of Experimental Marine Biology and Ecology 502, 12–25. 10.1016/j.jembe.2017.01.007

St. John, J., 2011. SeqPrep: Tool for stripping adaptors and/or merging paired reads with overlap into single reads.

Stoeck, T., Bass, D., Nebel, M., Christen, R., Jones, M., Breiner, H., Richards, T., 2010. Multiple marker parallel tag environmental DNA sequencing reveals a highly complex eukaryotic community in marine anoxic water. Molecular Ecology 19, 21–31. 10.1111/j.1365-294X.2009.04480.x

Sun, Z., Liu, J., Zhang, M., Wang, T., Huang, S., Weiss, S.T., Liu, Y.-Y., 2023. Removal of false positives in metagenomics-based taxonomy profiling via targeting Type IIB restriction sites. Nat Commun 14, 5321. 10.1038/s41467-023-41099-8

Thomas, A.C., Deagle, B.E., Eveson, J.P., Harsch, C.H., Trites, A.W., 2016. Quantitative DNA metabarcoding: improved estimates of species proportional biomass using correction factors derived from control material. Mol Ecol Resour 16, 714–726. 10.1111/1755-0998.12490

van Der Heijden, L., Niquil, N., Haraldsson, M., Asmus, R., Pacella, S., Graeve, M., Rzeznik-Orignac, J., Asmus, H., Saint-Beat, B., Lebreton, B., 2020. Quantitative food web modeling unravels the importance of the microphytobenthos-meiofauna pathway for a high trophic transfer by meiofauna in soft-bottom intertidal food webs. Ecological Modelling 430, 109129. 10.1016/j.ecolmodel.2020.109129

van der Loos, L., Nijland, R., 2021. Biases in bulk: DNA metabarcoding of marine communities and the methodology involved. Molecular Ecology 30, 3270–3288. 10.1111/mec.15592

Wilcox, T.M., Zarn, K.E., Piggott, M.P., Young, M.K., McKelvey, K.S., Schwartz, M.K., 2018. Capture enrichment of aquatic environmental DNA: A first proof of concept. Mol Ecol Resour 18, 1392–1401. 10.1111/1755-0998.12928

Ye, S.H., Siddle, K.J., Park, D.J., Sabeti, P.C., 2019. Benchmarking Metagenomics Tools for Taxonomic Classification. Cell 178, 779–794. 10.1016/j.cell.2019.07.010

Zhan, A., MacIsaac, H.J., 2015. Rare biosphere exploration using high-throughput sequencing: research progress and perspectives. Conserv Genet 16, 513–522. 10.1007/s10592-014-0678-9

