## Supplementary information for "Benchmarking the quantitative performance of metabarcoding and shotgun sequencing using mock communities of marine nematodes"

**Running title**: Metabarcoding vs. shotgun sequencing? Quantitative insights from mock communities

**Keywords:** Abundance quantification, biomonitoring, community ecology, meiofauna, organismal traits

### Supplementary Text

#### Text1 PCR amplification conditions of 18S and 28S metabarcoding

The first round of PCR using the 18S primers was performed in 25-μl reaction mixtures containing 12.5 μl of Q5HS master Mix (New England BioLabs), 0.5 μl of each primer (10 nM; with attached adapter sequence), 5 μl of template DNA, and 6.5 μl of PCR molecular biology grade water (Thermo Fisher Scientific). All reactions were performed in triplicate under the following thermal conditions: 98°C of initial denaturation for 30 s, followed by 15 cycles of 98°C denaturation for 10 s, 50°C annealing for 30 s, and 72°C elongation for 30 s, with a final extension step of 10 min at 72°C. First-round amplicons were pooled and then cleaned by treatment with 0.1 mL of exonuclease I (New England BioLabs) and 0.2 mL of thermosensitive alkaline phosphatase (Promega), with a 15-min incubation at 37°C followed by a 15-min incubation at 74°C to terminate the reaction. The second round of PCR was carried out using indexing primers in order to equip each sample with a unique combination of forward and reverse index sequences. The thermal conditions for this PCR were 3 min at 95°C, 15 cycles with 1 cycle consisting of 30 s at 95°C, 30 s at 55°C, and 30 s at 72°C, with a final elongation of 5 min at 72°C. The final PCR products (triplicate pooled in one tube) were then cleaned using Agencourt AMPure XP magnetic beads according to the manufacturer's instructions (Beckman Coulter). DNA concentrations were determined using a Qubit 2.0 fluorometer and the double-stranded DNA BR assay kit (Invitrogen) before the samples were standardized and pooled. Libraries were sequenced on an Illumina MiSeq V3 platform with 2 × 300-bp paired-end setup at the National Genomics Infrastructure (NGI) in Stockholm, Sweden (SciLifeLab, Stockholm, Sweden).

The library preparation procedures used for 18S metabarcoding were also applied to 28S metabarcoding, except that the thermal conditions of the first-round PCR were 60 s at 96°C, followed by 15 s at 96°C, 30 s at 58 °C, and 90 s at 72°C, with a final elongation of 10 min at 72°C. In addition, eight Nematoda were individually sequenced using barcoding and the generated sequences (Supplementary Table S2) were then used to create an in-house reference database of the 28S D3-D5 region for this study, owing to the lack of a sufficient database for 28S targeting marine nematodes.

### Supplementary Figures

**Figure S1** Maximum-likelihood phylogenetic tree of the 28S rRNA gene in-house reference sequences. The nematode genera are colored according to their hypoxia sensitivity: tolerant taxa in dark orange, less sensitive taxa in light orange, and sensitive taxa in very light orange. The 28S reference sequences were aligned using MAFFT v.7 with the L-INS-I local pair alignment method, and phylogenetic relationships were inferred using the maximum-likelihood method in IQ-TREE. The resulting tree was visualized in iTOL v. 7.5.1.

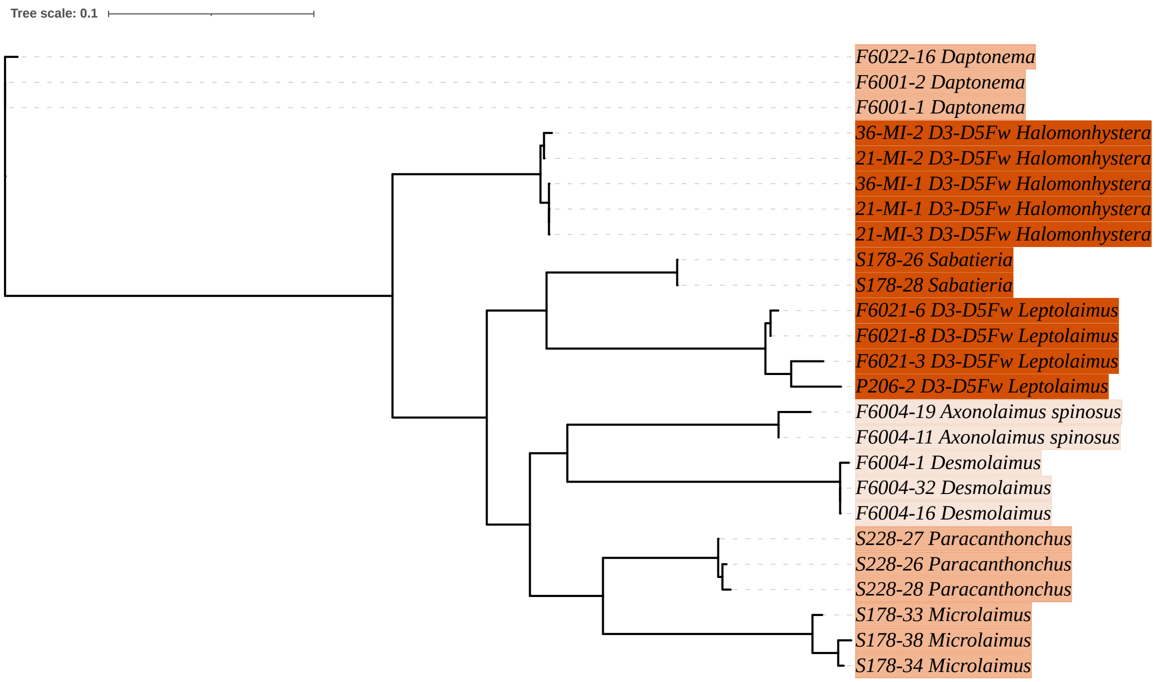

**Figure S2** Number of ASVs or contigs recovered across different nematode genera.

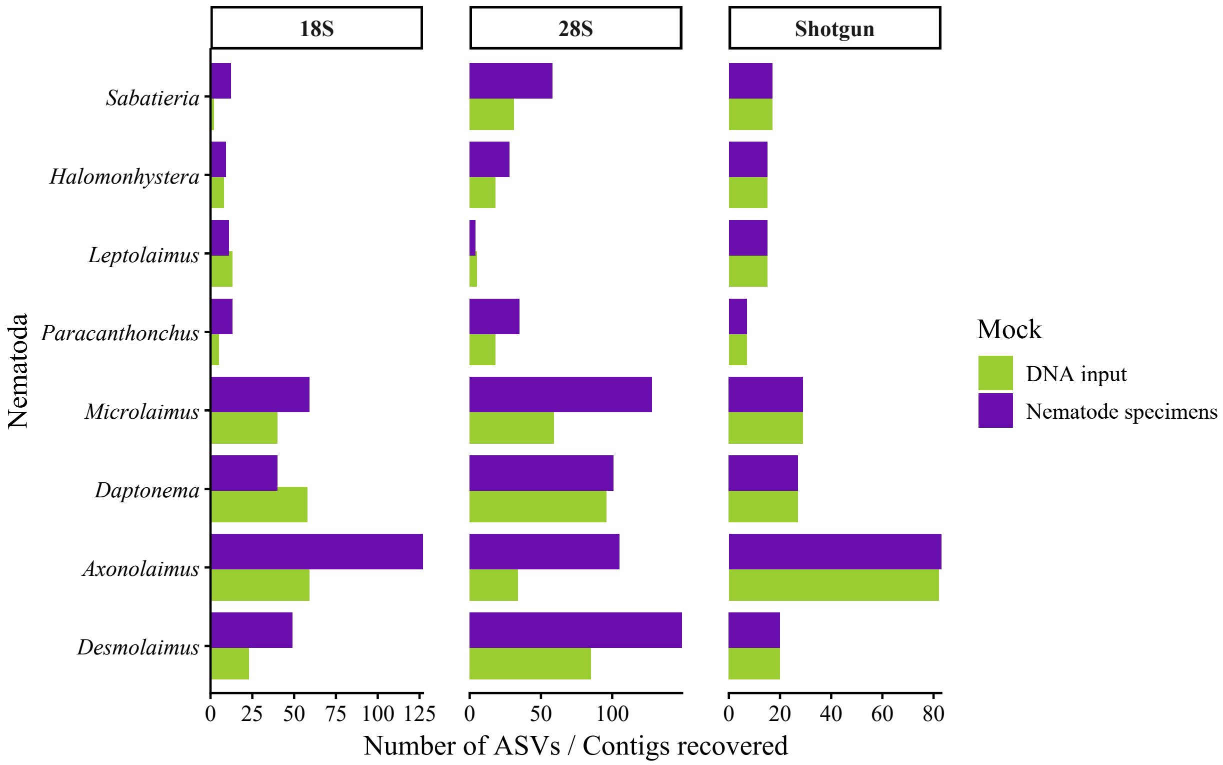

**Figure S3** The comparison of sequence reads that were identified as false-positives and non-target taxa in the 18S, 28S and shotgun datasets (those assigned to nematode outside of the target nematode genera as well as those assigned to non-nematodes). Significance differences at $P < 0.001$ determined in the corresponding Wilcoxon test are shown in the figure. The presented relative abundances were calculated by the proportion of those sequence reads in the respective datasets.

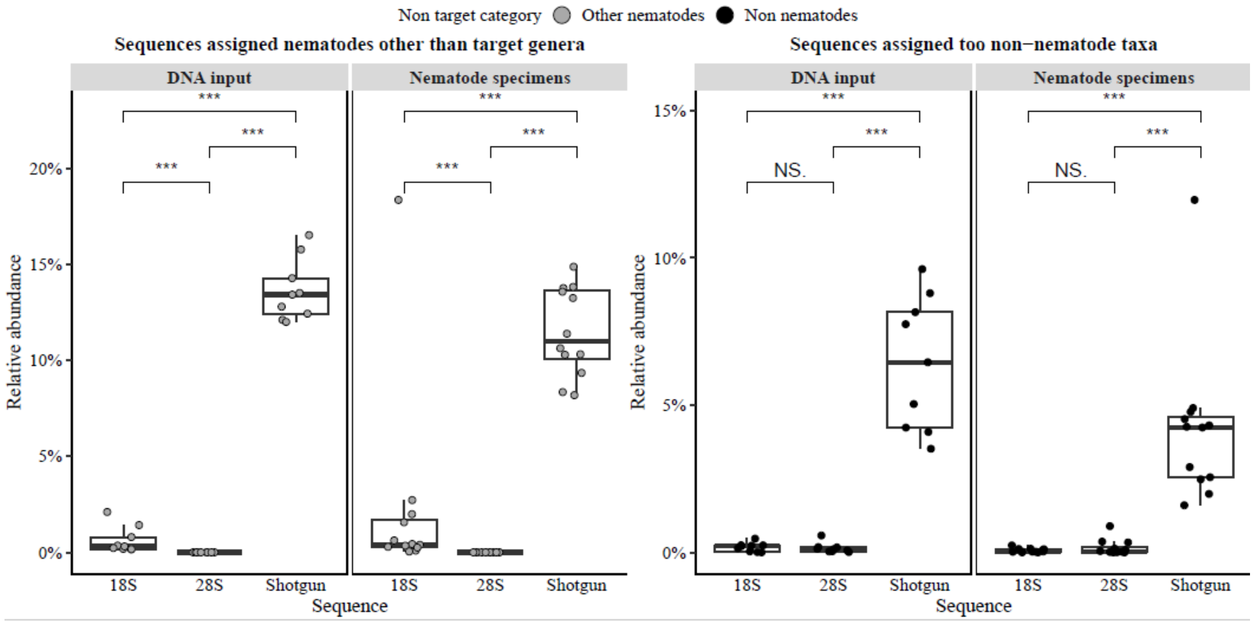

**Figure S4** Dissimilarity of the mock communities with an even or staggered composition. **A**) Principal coordinates analysis (PCoA) of the mock communities composed of inputted DNA. **B**) PCoA of the mock communities composed of nematode specimens. For both panels, PCo 1 and PCo2 represent the percentage of the community variation explained by the analysis.

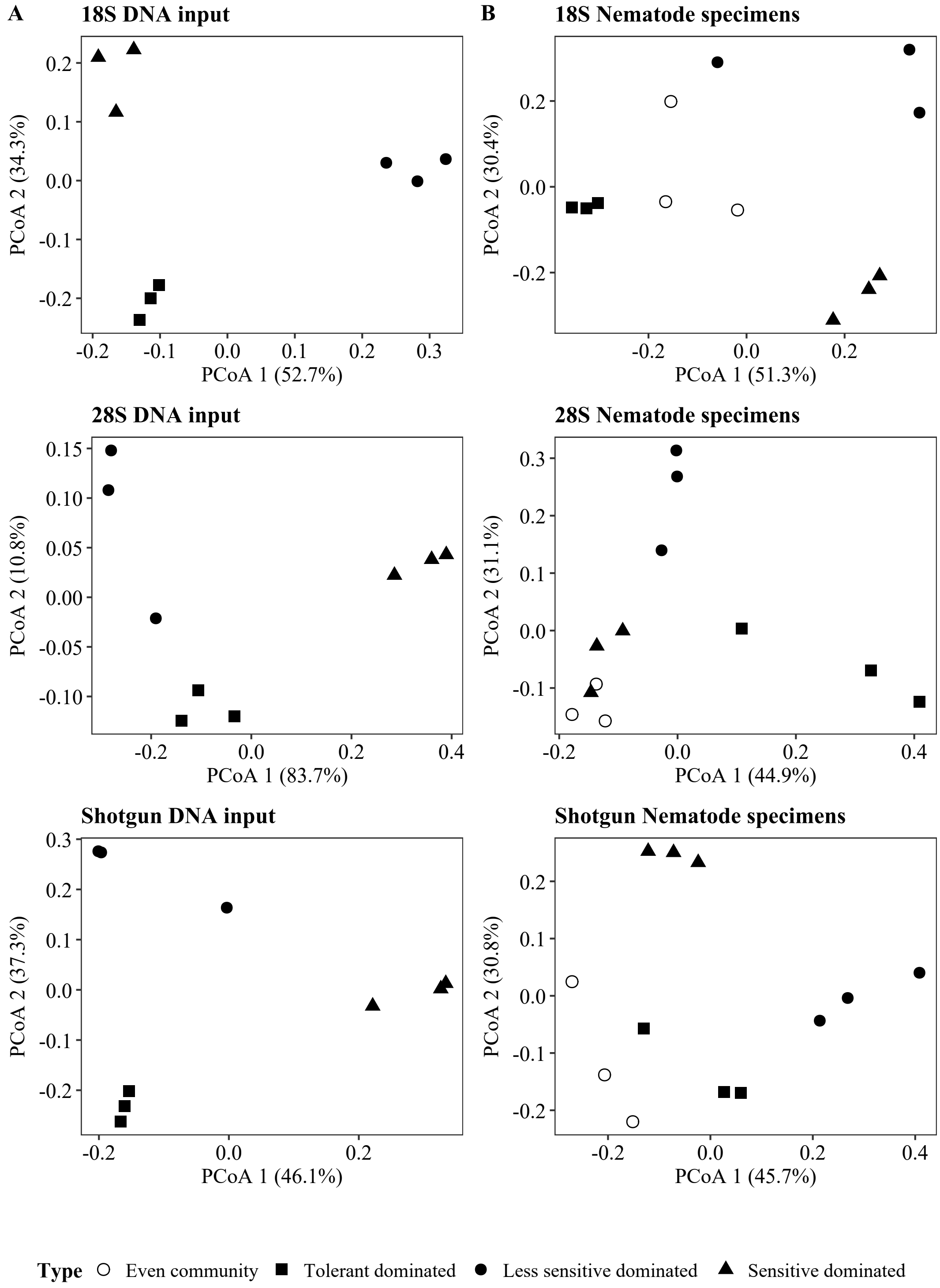

**Figure S5** Observed relative abundance obtained from high-throughput sequencing plotted against the expected proportion for each Nematoda genus in the mock communities composed of DNA input (**A–C, experiment 1**) and individual nematode specimens (**D–F, experiment 2**). The correlation coefficients and *P* value obtained from the Pearson correlation analysis are reported in each panel.

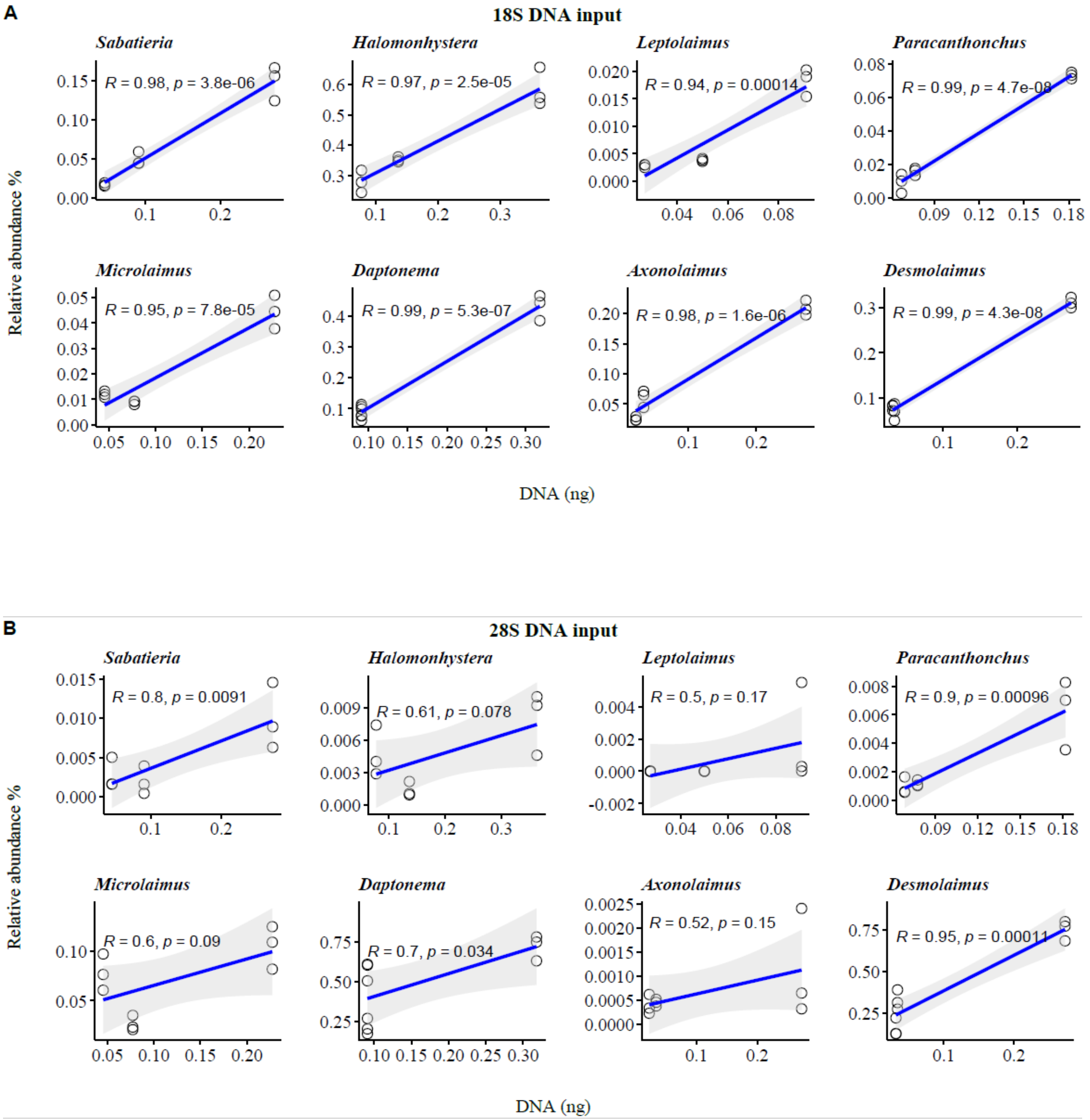

C

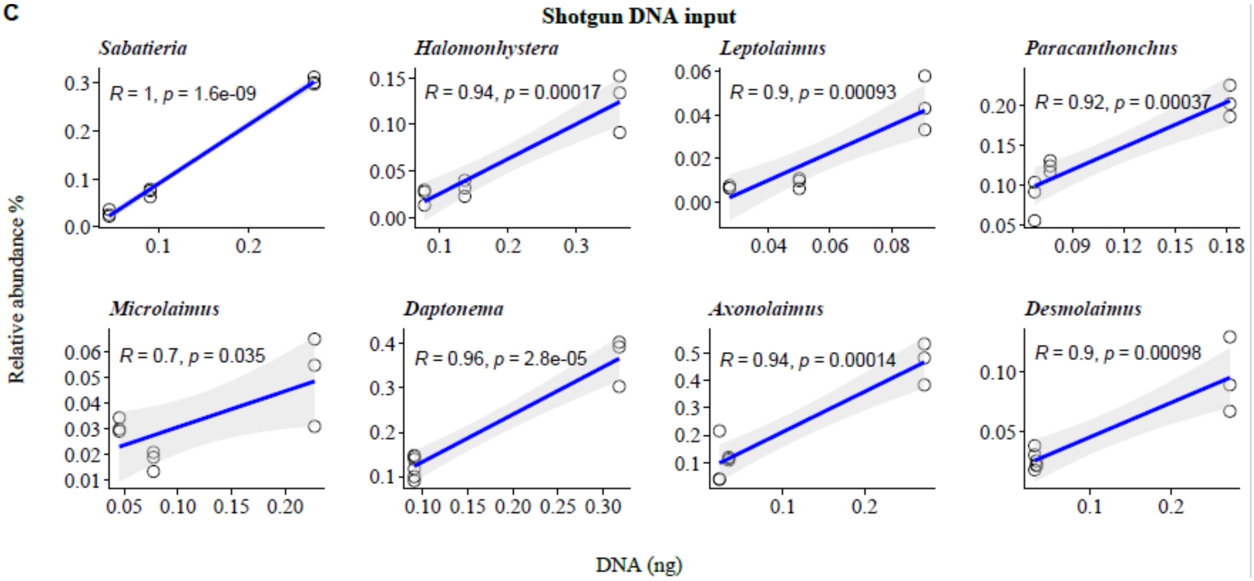
Shotgun DNA input
Sabatieria
R = 1, p = 1.6e-09
Halomonhystera
R = 0.94, p = 0.00017
Leptolaimus
R = 0.9, p = 0.00093
Paracanthonchus
R = 0.92, p = 0.00037
Microlaimus
R = 0.7, p = 0.035
Daptonema
R = 0.96, p = 2.8e-05
Axonolaimus
R = 0.94, p = 0.00014
Desmolaimus
R = 0.9, p = 0.00098
Relative abundance %
DNA (ng)

D

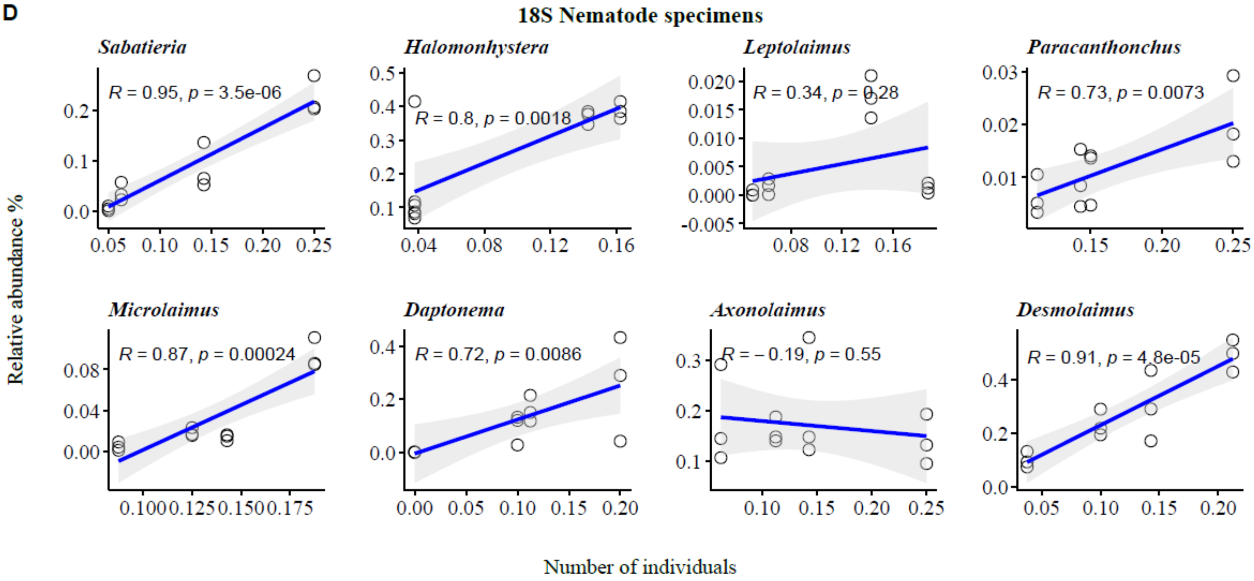
18S Nematode specimens
Sabatieria
R = 0.95, p = 3.5e-06
Halomonhystera
R = 0.8, p = 0.0018
Leptolaimus
R = 0.34, p = 0.28
Paracanthonchus
R = 0.73, p = 0.0073
Microlaimus
R = 0.87, p = 0.00024
Daptonema
R = 0.72, p = 0.0086
Axonolaimus
R = −0.19, p = 0.55
Desmolaimus
R = 0.91, p = 4.8e-05
Relative abundance %
Number of individuals

E

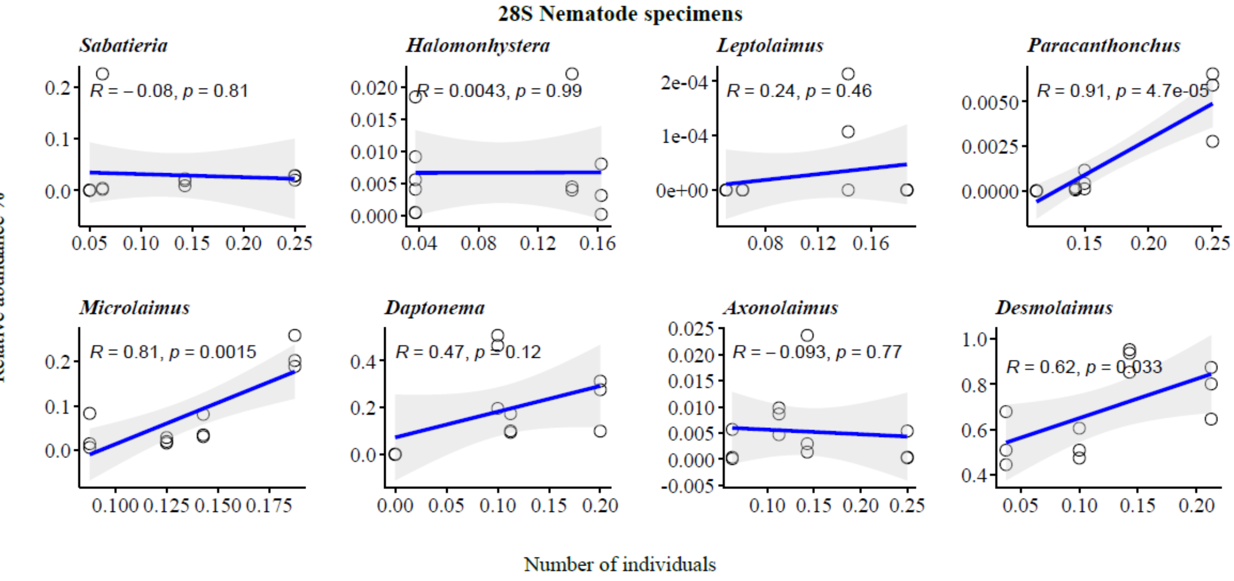
28S Nematode specimens
Sabatieria
R = −0.08, p = 0.81
Halomonhystera
R = 0.0043, p = 0.99
Leptolaimus
R = 0.24, p = 0.46
Paracanthonchus
R = 0.91, p = 4.7e-05
Microlaimus
R = 0.81, p = 0.0015
Daptonema
R = 0.47, p = 0.12
Axonolaimus
R = −0.093, p = 0.77
Desmolaimus
R = 0.62, p = 0.033
Relative abundance %
Number of individuals

F

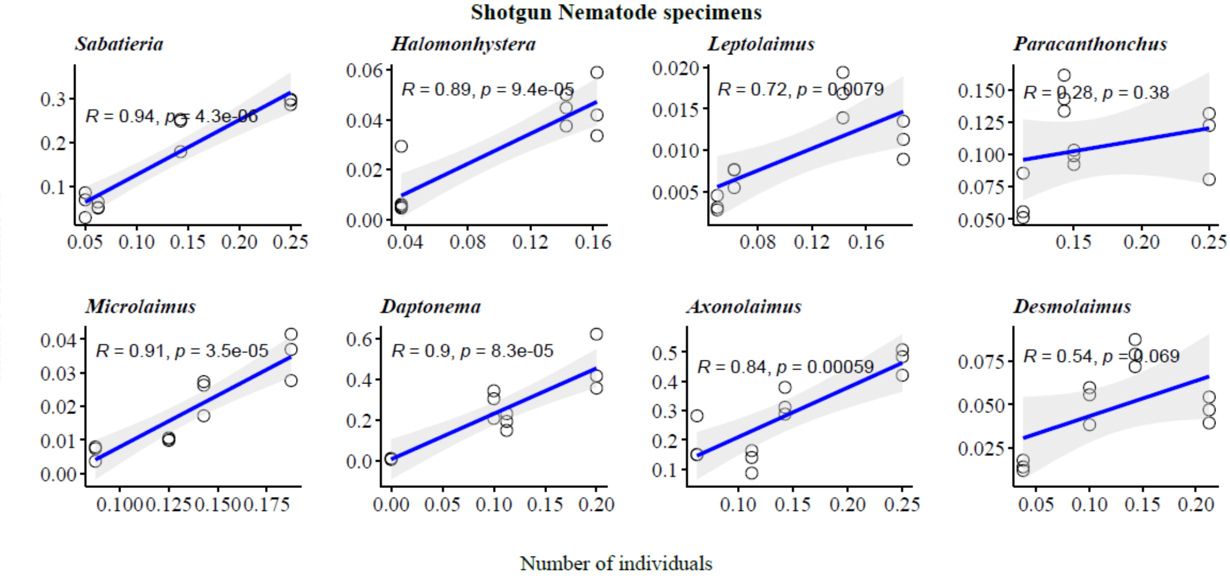
Shotgun Nematode specimens
Sabatieria
R = 0.94, p = 4.3e-06
Halomonhystera
R = 0.89, p = 9.4e-05
Leptolaimus
R = 0.72, p = 0.0079
Paracanthonchus
R = 0.28, p = 0.38
Microlaimus
R = 0.91, p = 3.5e-05
Daptonema
R = 0.9, p = 8.3e-05
Axonolaimus
R = 0.84, p = 0.00059
Desmolaimus
R = 0.54, p = 0.069
Relative abundance %
Number of individuals

**Figure S6** Bar-plots illustrating the relationship between categorical body-size groups and read abundance across the different sequencing approaches for each genus in the even mock communities. Because all genera were represented by the same number of individuals in the even mock communities, any effect of body size, if it acts as a rough proxy for biomass, could therefore be reflected in read abundance. The target nematode genera were categorized into four body sizes as follows: small (*Leptolaimus*, *Microlaimus*), intermediate slender-bodied (*Paracanthonchus*, *Desmolaimus*), intermediate thick-bodied (*Sabatieria*), large (*Halomonhystera*, *Axonolaimus*).

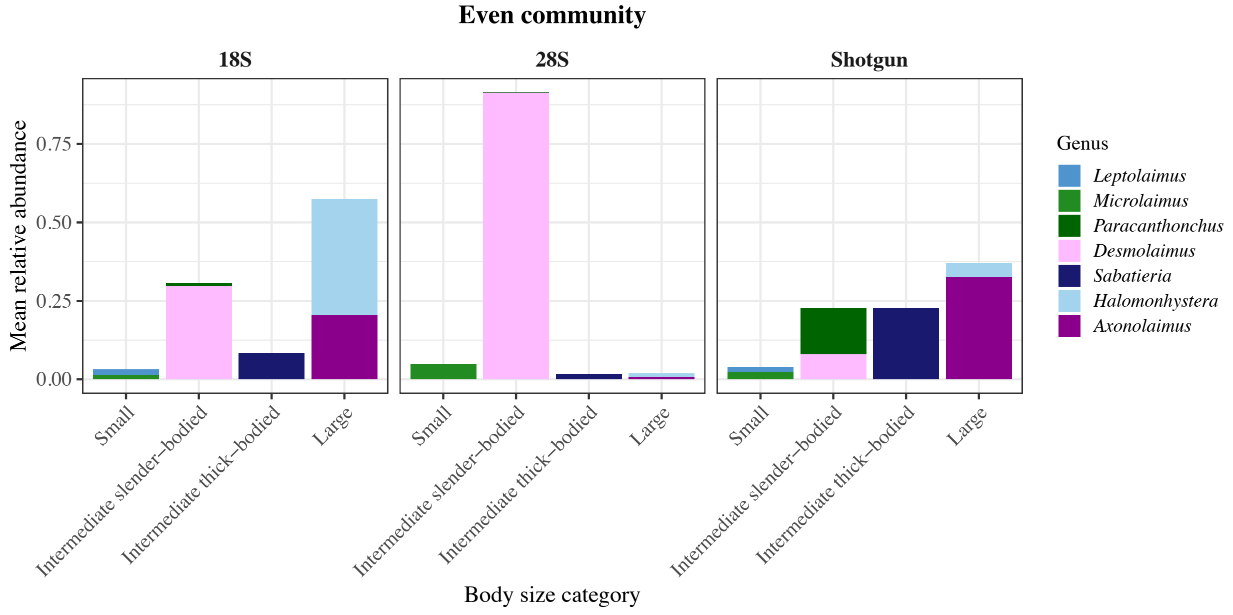

**Supplementary Tables (are presented in excel sheets)**

**Table S1** DNA quantity (ng) (**A**) and the number of individuals (**B**) for eight nematode genera used to construct the mock communities, with their corresponding expected proportions. In (**A**), values represent the DNA amount required for each genus in a single replicate community (three replicates in total). Although DNA input varied among genera within each mock community, the total DNA amount was 22 ng for all communities. In (B), values represent the number of specimens required for each genus in a single replicate community. *Daptonema* was excluded from the even mock community because of insufficient specimen availability.

**Table S2** Barcoding sequences retrieved from 28S D3-D5 of the eight Nematoda genera.

**Table S3** Sequence summary of the 18S metabarcoding results, including the number of sequences retained per ASV under each sample id, BLAST query results, and taxonomic classification. Non-target hits are highlighted in orange and comprise sequences assigned either to nematodes outside the target taxa (referred to as "Other nematodes" in the manuscript), or to non-nematodes. Sequences highlighted in gray either got no blast hit or did not have a top hit that passed filtering. The ASV IDs presented in this table refer to the sequence variants identified after DADA2 processing. Empty fields indicate that no assignment was available at that taxonomic level. Abbreviations of the BLAST output are as follows:

"sseqid" = subject sequence ID; the ID of the matching sequence in the database.
"staxids" = subject taxonomy IDs; the NCBI taxonomy ID(s) of the matched subject sequence.
"sscinames" = subject taxonomy name; the name corresponding to the NCBI taxonomy ID(s) of the matched subject sequence.
"salltitles" = all titles of the matched subject.
"pident" = percent identity; the percentage of identical matches between query and subject over the aligned region.
"length" = alignment length; the number of positions included in the alignment.
"mismatch" = number of mismatches; How many positions differ between query and subject in the alignment.
"gapopen" = number of gap openings; How many gaps were opened in the alignment.
"qstart" = query start; Start position of the alignment in your query sequence.
"qend" = query end; End position of the alignment in your query sequence.
"sstart" = subject start; Start position of the alignment in the database subject sequence.
"send" = subject end; End position of the alignment in the database subject sequence.
"evalue" = expectation value; A measure of how likely the match could happen by chance.
"bitscore" = bit score; A normalized alignment score.
"qlen" = Length of query sequence.
"qcovhsp" = query coverage per HSP; The percentage of your query sequence covered by the alignment to that HSP of a subject.

“qcovs” = query coverage per subject; The percentage of your query sequence covered by the alignment to that subject.

**Note:** The majority of ASVs showed relative abundances <0.5% per sample. An exception was observed for ASV 18S_11 (18.1% relative abundance), assigned to Oncholaimidae, detected only in sample 18S_Exp2_Sen1 in the 18S dataset. This taxon was not part of the mock community design and is therefore interpreted as a false positive, potentially resulting from accidental inclusion of a non-target specimen during sorting (>2,500 individuals processed). This isolated observation does not affect the overall pattern, as the expected sensitive-taxa-dominated community composition was consistently recovered in the other replicates.

**Table S4** Table S4 Sequence summary of the 28S metabarcoding results, including the number of sequences retained per ASV under each sample id, taxonomic classification, and BLAST query results. The ASV IDs presented in this table refer to the sequence variants identified after DADA2 processing. Non-target hits are highlighted in orange and comprise sequences assigned either to nematodes outside the target taxa (referred to as “Other nematodes” in the manuscript), or to non-nematodes. Sequences highlighted in gray either got no blast hit or did not have a top hit that passed filtering. Abbreviations of the BLAST output are as follows:
“sseqid” = subject sequence ID; the ID of the matching sequence in the database.
“staxids” = subject taxonomy IDs; the NCBI taxonomy ID(s) of the matched subject sequence.
“sscinames” = subject taxonomy name; the name corresponding to the NCBI taxonomy ID(s) of the matched subject sequence.
“salltitles” = all titles of the matched subject.
“pident” = percent identity; the percentage of identical matches between query and subject over the aligned region.
“length” = alignment length; the number of positions included in the alignment.
“mismatch” = number of mismatches; How many positions differ between query and subject in the alignment.
“gapopen” = number of gap openings; How many gaps were opened in the alignment.
“qstart” = query start; Start position of the alignment in your query sequence.
“qend” = query end; End position of the alignment in your query sequence.
“sstart” = subject start; Start position of the alignment in the database subject sequence.
“send” = subject end; End position of the alignment in the database subject sequence.
“evalue” = expectation value; A measure of how likely the match could happen by chance.
“bitscore” = bit score; A normalized alignment score.
“qlen” = Length of query sequence.
“qcovhsp” = query coverage per HSP; The percentage of your query sequence covered by the alignment to that HSP of a subject.
“qcovs” = query coverage per subject; The percentage of your query sequence covered by the alignment to that subject.

**Table S5** Table S5 Sequence summary of the shotgun results, including the number sequence counts per contig assembled under each sample id, BLAST query results, and the taxonomic classification. non-target hits are highlighted in orange and comprise sequences assigned either to nematodes outside the target taxa (referred to as "Other nematodes" in the manuscript), or to non-nematodes. Sequences highlighted in gray either got no blast hit or did not have a top hit that passed filtering.

**Table S6** Results of the Mantel test assessing differences among mock communities sequenced using 18S metabarcoding, 28S metabarcoding, and shotgun sequencing.

**Table S7** The coefficients and *P*-values obtained from a Pearson correlation analysis of the relative abundance vs. the expected proportion of the total DNA quantity in the mock communities. 'Type' represents the mock communities constructed using DNA input or individual nematode specimens. 'Sequences' indicates that the analyzed sequences were retrieved from 18S metabarcoding, 28S metabarcoding, or shotgun sequencing. '95%low' and '95%upp' represent the lower and upper limits of the 95% confidence intervals of the coefficient, respectively. 'SE' represents the standard errors calculated from the lower and upper limits of the 95% confidence intervals. P-values in bold indicate a correlation significance < 0.05.

**Table S8** Recovery of number of genera in the different mock communities, the number of missing by mock community and genus.
